# Estimating scale-specific and localized spatial patterns in allele frequency

**DOI:** 10.1101/2022.03.21.485229

**Authors:** Jesse R. Lasky, Margarita Takou, Diana Gamba, Timothy H. Keitt

## Abstract

Characterizing spatial patterns in allele frequencies is fundamental to evolutionary biology because these patterns contain evidence of underlying processes. However, the spatial scales at which gene flow, changing selection, and drift act are often unknown. Many of these processes can operate inconsistently across space, causing non-stationary patterns. We present a wavelet approach to characterize spatial pattern in allele frequency that helps solve these problems. We show how our approach can characterize spatial patterns in relatedness at multiple spatial scales, i.e. a multi-locus wavelet genetic dissimilarity. We also develop wavelet tests of spatial differentiation in allele frequency and quantitative trait loci (QTL). With simulation we illustrate these methods under different scenarios. We also apply our approach to natural populations of *Arabidopsis thaliana* to characterize population structure and identify locally-adapted loci across scales. We find, for example, that Arabidopsis flowering time QTL show significantly elevated genetic differentiation at 300 to 1300 km scales. Wavelet transforms of allele frequencies offer a flexible way to reveal geographic patterns and underlying evolutionary processes.

## 1 Introduction

Geographic clines in allele frequency are a classic pattern in evolutionary biology, being frequently observed in nature and having extensive theory for the underlying processes. For example, theory describes how limited gene flow and drift (Wright 1931) or changing selection (Haldane 1948) can generate allele frequency differences between populations. Accordingly, researchers often estimate and model spatial allele frequency patterns to make inferences about underlying evolutionary and ecological mechanisms. To do so, researchers often divide sampled individuals into discrete groups (populations) among which differences in allele frequencies are calculated. A common such approach involves estimating *F_ST_*, the proportion of total allele frequency variation that differs between discrete populations (Lewontin and Krakauer 1973; Wright 1949).

However, many species exist as more or less continuously distributed populations. Theoretical study of allele frequency change across continuous populations began as early as Wright (1943) and Maĺecot (1948), who found expectations for genetic differentiation or kinship as functions of gene flow and geographic distance. Later progress included diffusion models (Nagylaki 1978) and stepping stone/lattice models (Kimura and Weiss 1964) giving expectations for correlation in allele frequencies across distance, and models accounting for population regulation by negative density dependence (Nick H. Barton, Depaulis, and Etheridge 2002).

Despite these theoretical advances, the statistical tools for inference on continuously distributed populations have lagged (Bradburd and Peter L. Ralph 2019; Hancock, Toczydlowski, and Bradburd 2023). Nevertheless, statistical approaches to studying spatial pattern in continuous populations include models relating landscape features to gene flow (McRae et al. 2008), calculating correlations between spatial functions and genotype (Wagner, Chávez-Pesqueira, and Forester 2017; Yang et al. 2012), and applying discrete landscape grids to identify geographic regions where genetic turnover is particularly high or low (Petkova, Novembre, and Stephens 2016). Approaches have been developed to estimate the average distance of gene flow from the slope of genetic divergence versus geographic distance (Rousset 2000; X. Vekemans and O. J. Hardy 2004), to estimate localized genetic “neighborhoods” (Shirk and Cushman 2014; Wright 1946), and to model both discrete and continuous relatedness patterns simultaneously (Bradburd, G. M. Coop, and Peter L. Ralph 2018).

In recent years researchers have collected many large, broadly distributed DNA sequence datasets from diverse species (Alonso-Blanco et al. 2016; Machado et al. 2021; J. Wang et al. 2020; Yeaman et al. 2016). Statistical inference can be applied to these data to understand gene flow, demographic histories, and spatially-varying selection. Despite the progress made by previous approaches, there remain challenges.

### 1.1 The form and scale of relevant spatial patterns is unknown

Humans can infer seemingly meaningful patterns in even randomly generated images (Ayton and Fischer 2004; Blakemore et al. 2003; Fyfe et al. 2008). So what are the spatial patterns we are looking for? The functional forms (i.e. shapes) of both spatially-varying selection and neutral processes (e.g. dispersal kernels) are often unknown, as are the forms of resulting spatial patterns. For example, the specific environmental gradients driving changing selection are often not known, nor is the spatial scale at which they act, and whether they change at the same rate consistently across a landscape.

In the case of neutral processes, a homogeneous landscape approximately at equilibrium is rarely of interest to empiricists. Instead, the influence of heterogeneous landscapes (Manel et al. 2003) and historical contingency is usually a major force behind spatial patterns in allele frequency and traits (Excoffier and Ray 2008). As a result, researchers often attempt to characterize spatial patterns of relatedness and genetic similarity to make inferences about variation in gene flow (McRae et al. 2008; Peterman 2018; I. J. Wang, Savage, and Bradley Shaffer 2009) and recent population expansion (Slatkin 1993). The influence of gene flow, drift, and range expansion can occur at a variety of spatial scales, and in different ways across a heterogenous landscape. For example, the rate at which relatedness decays over geographic distance can change abruptly at major barriers (Rosenberg et al. 2005). However, the scale-specificity and non-stationarity of such patterns can be challenging to characterize.

### 1.2 The spatially-varying selective gradients causing local adaptation are unknown

One important force behind allele frequency clines is changing selection due to environmental gradients, resulting in local adaptation. However, it is often not clear what environmental gradients drive local adaptation (Kawecki and Ebert 2004). This is especially true of non-model systems and those with little existing natural history knowledge. Even for well-studied species, it is not trivial to identify the specific environmental conditions that change in space and drive local adaptation. Ecology is complex, and abiotic and biotic conditions are highdimensional. Rather than *a priori* selection of a putative selective gradient, an alternative approach is to search for spatial patterns in allele frequencies that cannot be explained by neutral processes. This approach is embodied by several statistics and approaches, such as *F_ST_* (Weir and Cockerham 1984), *XtX* (Gautier 2015), spatial ancestry analysis (SPA) (Yang et al. 2012), Moran’s eigenvector maps (MEMs) (Wagner, Chávez-Pesqueira, and Forester 2017), and others.

### 1.3 Many approaches rely on discretization of population boundaries

Some of the aforementioned approaches rely on dividing sampled individuals into discrete spatial groups. *F_ST_* is one such approach, that was introduced by Wright (1949) and defined as the “correlation between random gametes, drawn from the same subpopulation, relative to the total”, where the definition of “total” has been interpreted differently by different authors (Bhatia et al. 2013). The classic approach of calculating *F_ST_* to test for selection was usually applied to a small number of locations, a situation when discretization (i.e. deciding which individuals genotyped belong in which population) was a simpler problem. Current studies often sample and sequence individuals from hundreds of locations, and so the best approach for discretizing these genotyped individuals into defined ‘populations’ is less clear. In addition to the spatial scale of subpopulations, at issue is precisely where to place the boundaries between populations. The problem is enhanced for broadly distributed species, connected by gene flow, that lack clear spatially distinct populations (Emily B. Josephs et al. 2019). Even if clustering algorithms appear to show clustering of genotypes, these methods can be sensitive to sampling bias (e.g. geographic clustering) and can mislead as to the existence of discrete subpopulations (Frantz et al. 2009; Serre and Pääbo 2004).

Some approaches are not limited by discretization, and might be generally termed “population-agnostic” because discrete populations are not defined. These instead use ordination of genetic loci or geographic location. Approaches that use ordination (such as PCA) of genetic loci look for particular loci with strong loadings on PCs (Duforet-Frebourg et al. 2016) or traits with an unexpectedly high correlation with individual PCs (Emily B. Josephs et al. 2019). Alternatively, ordination of distance or spatial neighborhood matrices can create spatial functions that can be used in correlation tests with genetic loci (Wagner, Chávez-Pesqueira, and Forester 2017). However, ordinations to create individual rotated axes are not done with respect to biology and so might not be ideal for characterizing biological patterns. For example, ordinations of genetic loci are heavily influenced by global outliers of genetic divergence (Peter, Petkova, and Novembre 2020) and uneven sampling (McVean 2009). Ordinations like PCA also often lack parametric null distributions for hypothesis testing.

### 1.4 Wavelet characterization of spatial pattern

Instead of discretizing sampled locations into populations, one could model allele frequencies with flexible but smooth functions. Wavelet transforms allow one to characterize the location and the scale or frequency of a signal (Daubechies 1992). Daubechies (1992) gives a nice analogy of wavelet transforms: they are akin to written music, which indicates a signal of a particular frequency (musical notes of different pitch) at a particular location (the time at which the note is played, in the case of music). Applying this analogy to genetics, the frequency is the rate at which allele frequencies change in space, and the location is the part of a landscape where allele frequencies change at this rate. Applying wavelet basis functions to spatial genetic data could allow us to characterize localized patterns in allele frequency, and dilating the scale of these functions could allow us to characterize scale-specific patterns in allele frequency (see Figure S1 for an example). Note that wavelets are distinct from Fourier analysis. Wavelets capture localized signals because the basis functions’ variance goes to zero moving away from the focal location, while Fourier can only capture global average patterns as it uses stationary (unchanging) basis functions. Wavelet transforms have had some recent applications in modeling ancestry along the genome (Groh and G. Coop 2023; Pugach et al. 2011) but have not been implemented to model geographic genetic patterns.

Keitt (2007) created a wavelet approach for characterizing spatial patterns in ecological communities. He used this approach to identify locations and scales with particular high community turnover, and created null-hypothesis testing of these patterns. These spatial patterns in the abundance of multiple species are closely analogous to spatial patterns in allele frequency of many genetic markers across the genome, and previous spatial genetic studies have also profited by borrowing tools from spatial community ecology (Fitzpatrick and Keller 2015; Jesse R. Lasky, Des Marais, et al. 2012). Here we modify and build on this approach to characterize spatial pattern in allele frequency across the genome and at individual loci.

## 2 Results

### 2.1 Wavelet characterization of spatial pattern in allele frequency

Our implementation here begins by following the work of Keitt (2007) in characterizing spatial community turnover, except that we characterize genomic patterns using allele frequencies of multiple loci in place of abundances of multiple species in ecological communities. In later sections of this paper we build off this approach and develop new tests for selection on specific loci. Our implementation of wavelets allows estimation of scale-specific signals (here, allele frequency clines) centered on a given point, *a, b*, in two-dimensional space. We use a version of the Difference-of-Gaussians (DoG) wavelet function (Figure S1) (Muraki 1995). We start with a Gaussian smoothing function centered at *a, b* for a set of sampling points Ω = *{*(*u*_1_*, v*_1_), (*u*_2_*, v*_2_), … (*u_n_, v_n_*)*}*, which takes the form

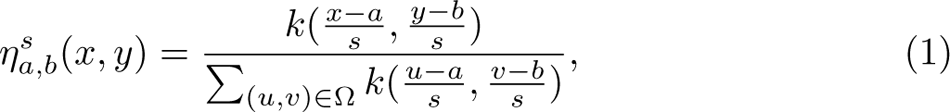

where *s* controls the scale of analysis and *k*(*x, y*) is the Gaussian kernel *k*(*x, y*) = *e^−^*^(^*^x^*^2 +^*^y^*^2)^*^/^*^2^. The DoG wavelet function then takes the form

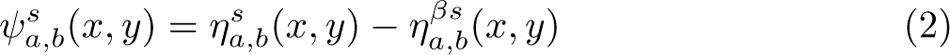

where *β >* 1, and so the larger scale smooth function is subtracted from the smaller scale smooth to characterize the scale-specific pattern. If we use *β* = 1.87, then the dominant scale of analysis resulting from the DoG is *s* distance units (Keitt 2007). This formulation of the wavelet kernel is similar in shape to the derivative-of-Gaussian kernel and has the advantage of maintaining admissibility (Daubechies 1992) even near boundaries, as each of the smoothing kernels *η^s^* are normalized over the samples such that their difference integrates to zero.

Let *f_i_*(*u, v*) be the major allele frequency of the *i*th locus from a set of *I* biallelic markers at a location with spatial coordinates *u, v*. The adaptive wavelet transform of allele frequency data at locus *i*, centered at *a, b* and at scale *s* is then

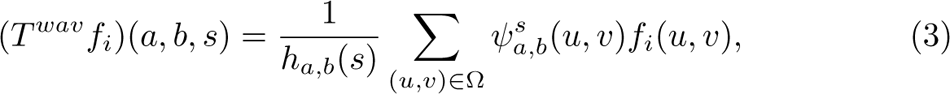

where the right summation is of the product of the smooth function and the allele frequencies across locations. The magnitude of this summation will be greatest when the DoG wavelet filter matches the allele frequency cline. That is, when the shape of the wavelet filter matches the allele frequency cline in space, the product of *ψ^s^* (*u, v*) and *f_i_*(*u, v*) will resonate (increase in amplitude) yielding greater variation among locations in (*T^wav^f_i_*)(*a, b, s*), the wavelet-transformed allele frequencies. When the spatial pattern in the wavelet filter and allele frequencies are discordant, the variation in their product, and hence the wavelet-transformed allele frequency, is reduced. For consistency, here we choose major allele frequency for *f_i_*(*u, v*), though in practice the signing of alleles has little impact on our results.

The *h_a,b_*(*s*) term in equation 3 is used to normalize the variation in the wavelet function so that the wavelet transforms *T^wav^f_i_*are comparable for different scales *s* and locations *a, b*:

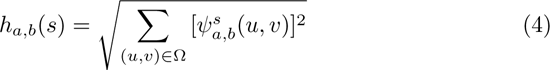

When *a, b* is far from locations in Ω relative to the scale *s*, the Gaussian functions [*η^s^* (*x, y*)] that make up the wavelet function *ψ* are only evaluated over a range where they remain close to zero. Thus unsampled geographic regions will have very small *h_a,b_*(*s*), the term used to normalize for local variation in the wavelet basis functions. In turn, very small *h_a,b_*(*s*) dramatically and undesirably inflates the wavelet transformed allele frequencies (equation 3) in these geographic regions where there is little sampling relative to *s*. For this reason we do not calculate the wavelet transform for locations *a, b* where there are no locations sampled closer than 2*s* distance units.

Below we illustrate how to apply this wavelet transform (equation 3) of spatial allele frequency patterns to characterize genome-wide patterns, as well as to test for local adaption at individual loci.

#### 2.1.1 Wavelet characterization of spatial pattern in multiple loci

Researchers are often interested in characterizing spatial patterns aggregated across multiple loci across the genome to understand patterns of relatedness, population structure, and demographic history. Here, we specifically want to characterize heterogeneity in spatial patterns, because this heterogeneity in pattern may reflect heterogeneity in underlying processes: where there is heterogeneity in migration rates, such as where there are migration barriers (Petkova, Novembre, and Stephens 2016), or where there are recent range expansions such that spatial patterns are farther from equilibrium (Slatkin 1993).

We use

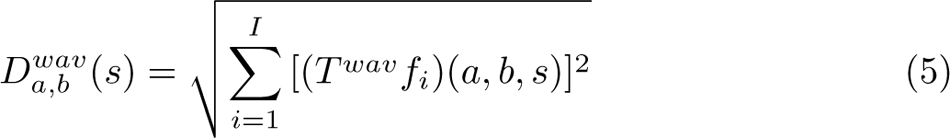

to calculate a “wavelet genetic distance” or “wavelet genetic dissimilarity.” This wavelet genetic dissimilarity is computed as the Euclidean distance (in the space of multiple loci’s allele frequencies) between the genetic composition centered at *a, b* and other locations across *s* distance units. This wavelet genetic dissimilarity *D^wav^*(*s*) is localized in space and scale-specific. This quantity captures the level of genetic turnover at scale *s* centered at *a, b*, and is capturing similar information as the increase in average genetic distance between a genotype at *a, b* and other genotypes *s* distance units away. To obtain the average dissimilarity across the landscape, one can also calculate the mean of *D^wav^*(*s*) across locations *a, b* at each sampled site, to get a mean wavelet genetic dissimilarity for *s*. A benefit of using the wavelet transformation over sliding window approaches (e.g. Bishop, Chambers, and I. J. Wang 2023) is that wavelets smoothly incorporate patterns from samples that are not precisely *s* distance units away and can be centered at any location of the analyst’s choosing.

#### 2.1.2 Testing the null hypothesis of no spatial pattern in allele frequency

A null hypothesis of no spatial pattern in allele frequencies can be generated by permuting the location of sampled populations among each other. Most empirical systems are not panmictic, and so this null model is trivial in a sense. However, comparison with this null across scales and locations can reveal when systems shift from small-scale homogeneity (from local gene flow) to larger scale heterogeneity (from limited gene flow) (Keitt 2007).

#### 2.1.3 Simulated neutral patterns across a continuous landscape

To demonstrate the wavelet transformation of allele frequencies, and wavelet genetic dissimilarity function, we applied these tools to several simulated scenarios. First, we conducted forward landscape genetic simulations under neutrality using the SLiM software (Haller and Messer 2019), building off published approaches (C J Battey, Peter L Ralph, and Kern 2020). We simulated out-crossing, iteroparous, hermaphroditic organisms, with modest lifespans (average of *∼* 4 time steps). Individual fecundity was Poisson distributed, mating probability (determining paternity) was determined based on a Gaussian kernel (truncated at three standard deviations), and dispersal distance from mother was also Gaussian (C. Battey, Peter L Ralph, and Kern 2020). Individuals became mature in the time step following their dispersal. These parameters roughly approximate a short lived perennial plant with gene flow via pollen movement and seed dispersal. Competition reduced survival and decayed with distance following a Gaussian (truncated at three standard deviations, C J Battey, Peter L Ralph, and Kern 2020). Near landscape boundaries, survival was reduced to compensate for lower competition from beyond the landscape margin (C J Battey, Peter L Ralph, and Kern 2020). Code is available at GitHub (https://github.com/jesserlasky/WaveletSpatialGenetic).

We began by characterizing a simple scenario across a continuous landscape. We simulated a square two dimensional landscape measuring 25 units on each side. The standard deviation of mating and dispersal distance *σ* were both 0.2, yielding a combined standard deviation of gene flow distances of 0.24 [(3*σ*^2^*/*2)^1*/*2^]. In this first simulation there was no selection. The population was allowed to evolve for 100,000 time steps before we randomly sampled 200 individuals and 1,000 SNPs with a minor allele frequency of at least 0.05. The first two principal components (PCs) of these SNPs show smooth population structure across the landscape, and that these two PCs predict the spatial location of each sample (Figure S2).

To facilitate interpretation of wavelet transformed allele frequencies (*T^wav^f_i_*)(*a, b, s*) we provide two example loci *i* with distinct spatial patterns (Figure 1). The first locus has the greatest variance in wavelet transformed allele frequencies among sampled loci at *s* = 0.4 (Figure 1A-C) while the second locus has the greatest variance at *s* = 12.2 (Figure 1D-F).

**Figure 1:**
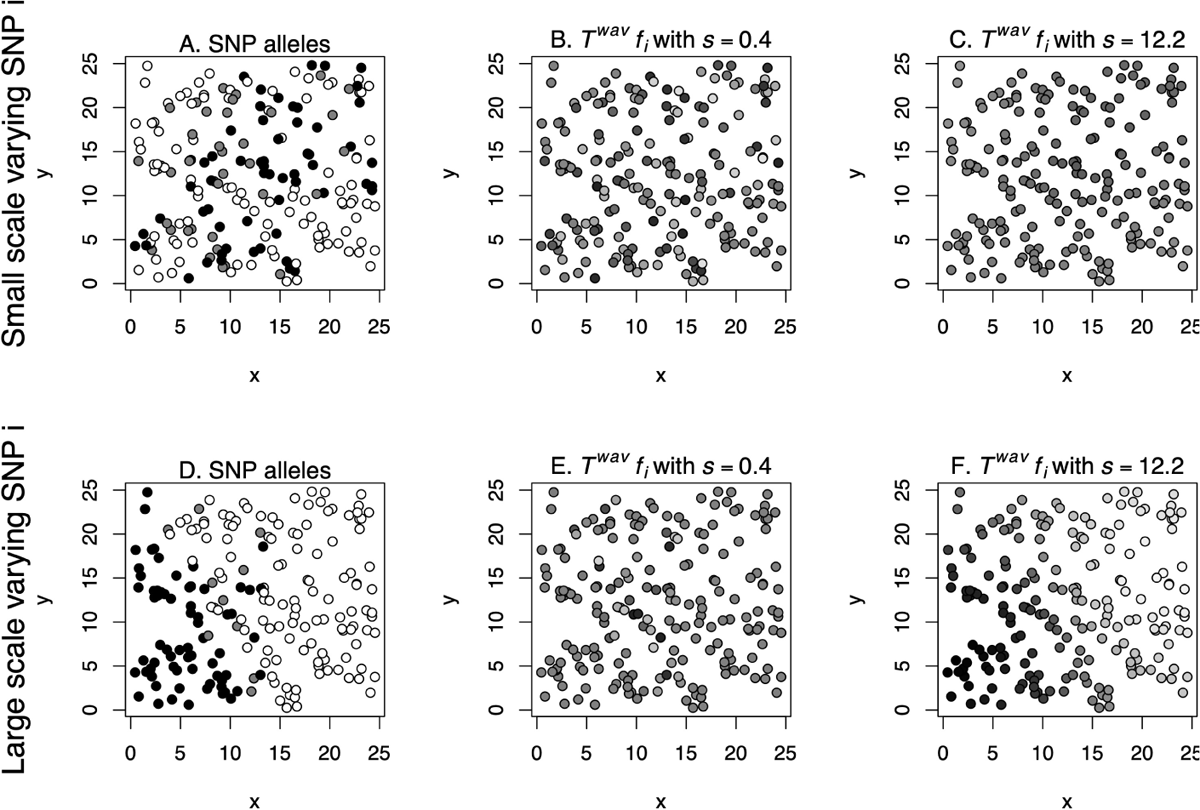
Two example SNPs (rows) with distinct spatial patterns. Shading shows either allelic variation (untransformed, A, D) or variation in wavelet transformed allele frequencies (*T^wav^f_i_*)(*a, b, s*) (B,C,E,F). The first locus (A-C) has the greatest variance in wavelet transformed allele frequency among sampled loci at *s* = 0.4. The second locus (D-F) has the greatest variance in wavelet transformed allele frequency at *s* = 12.2. For the SNP in the top row, the variance among locations in (*T^wav^f_i_*)(*a, b, s*) for *s* = 0.4 is 0.56 (visualized as shading in B), while it is only 0.17 for the SNP in the bottom row (E). For the SNP in the bottom row, the variance among locations in (*T^wav^f_i_*)(*a, b, s*) for *s* = 12.2 is 44.46 (visualized as shading in F), while it is only 1.24 for the SNP in the top row (C).

We then calculated wavelet dissimilarity *D^wav^*(*s*), aggregating the signals in (*T^wav^f_i_*)(*a, b, s*) across loci *i*, for each sampled location at a range of spatial scales *s*. Here and below we use a set of scales increasing by a constant log distance interval, as genetic distances are often linearly correlated to log geographic distances in two dimensions (F. Rousset 1997). The mean across sampled locations for each scale was calculated and compared to the null distribution for that scale (Figure S2). The null was generated by permuting locations of sampled individuals as described above, and observed mean of dissimilarity was considered significant if it was below the 2.5 percentile or above the 97.5 percentile of dissimilarity from null permutations.

When comparing our simulated data to the null, we found that mean wavelet genetic dissimilarity was significantly less than expected under the null model at scales *s ≤* 0.93, due to local homogenization by gene flow (standard deviation = 0.24). At scales *s ≥* 1.24, wavelet dissimilarity was significantly greater than expected, due to isolation by distance, with monotonically increasing wavelet genetic dissimilarity at greater scales (Figure S2).

To demonstrate how the scale of gene flow influences the wavelet dissimilarity *D^wav^*(*s*), we also conducted identical simulations as described above but instead with standard deviations of mating and dispersal distances, *σ*, of 0.5, 1, 2, or 5, yielding combined standard deviations of gene flow distances of 0.61, 1.22, 2.45, and 6.12.

To verify that simulations were generating results consistent with theoretical expectations of continuous populations at equilibrium, we compared the simulated gene flow parameters with estimations from the simulated data based on theory. The slope of genetic differentiation versus geographic distance in two dimensions is expected to be proportional to the inverse of Wright’s neighborhood size, 4*πDσ*^2^, where *D* is the effective population density and *σ* is the standard deviation of gene flow (Rousset 2000; X. Vekemans and O. J. Hardy 2004; Wright 1943, 1946).

We estimated *D* using *N_e_* = (4*N −* 2)*/*(*V* + 2) where *N* is census population size and *V* is variance in lifetime reproductive output (Kimura and Crow 1963). We then divided this *N_e_* by landscape area (assuming evenly distribution across the landscape) to get effective density *D* (X. Vekemans and O. J. Hardy 2004). We used three different genetic distance or kinship metrics (Loiselle et al. 1995; Ritland 1996; Rousset 2000) and used the SPAGeDi v1.5 software (Olivier J. Hardy and Xavier Vekemans 2002) to estimate gene flow across a range of true gene flow parameters. Simulations were run for 100,000 time steps with parameters as described above. We calculated *V* using the lifetime reproductive output of the individuals dying in the last 50 time steps.

We found that the gene flow estimated using the slope of genetic versus geographic distance and *D* was closely matched by the simulation parameter value, especially for the Rousset (2000) genetic differentiation estimator (Figure S3). This matching suggests these simulations corresponded well with theory for continuous populations at equilibrium, despite ignoring the effects of negative density dependence, uneven distribution of individuals, and boundary effects (Nick H. Barton, Depaulis, and Etheridge 2002).

With increasing scale of gene flow we see a flatter change in wavelet dissimilarity across spatial scales (Figure S4). When gene flow is local, wavelet dissimilarity is low at small scales and high at large scales. At the large gene flow scale, the observed wavelet dissimilarity is indistinguishable from the panmictic null. We also ran the same analyses but using biased sampling along the landscape’s y-axis, so that 3/4 of samples were in the upper half of the landscape. Even with this bias, the wavelet dissimilarities across scales and gene flow parameters were essentially unchanged (Figure S5). To investigate sensitivity to landscape size, we also ran these same simulations with landscapes four times as large (50×50) and found similar patterns of wavelet dissimilarity across scales and simulated gene flows (Figure S6).

#### 2.1.4 Simulated long-term neutral patterns in a heterogeneous landscape

To assess if our approach could identify localized and scale-specific patterns of isolation by distance, we next simulated multiple scenarios where we expected spatial heterogeneity. First, we simulated neutral evolution across a simulated patchy landscape (generated from earlier work) (Jesse R. Lasky and Keitt 2013). This landscape contained a substantial portion of unsuitable habitat where arriving propagules perished. We used the same population parameters as previously and simulated 100,000 time steps to reach approximately stable relatedness patterns. We then calculated wavelet dissimilarity using 1,000 random SNPs of 200 sampled individuals.

Additionally, we sought to compare wavelet dissimilarities to more familiar metrics. To do so, we calculated pairwise euclidean genetic and geographic distances between samples, and did this for different subsets of samples and regions, so as to compare localized patterns in wavelet dissimilarity to localized patterns in pairwise distances.

In our landscape, wavelet dissimilarity showed localized and scale-specific patterns of low and high dissimilarity (Figure 2). Notably, the same two islands (top left and bottom right of landscape in Figure 2) have lower dissimilarity than expected at small scales and are more dissimilar than expected at larger scales. Stated another way, these islands have low diversity locally (e.g. within populations), as can be seen by the slow increase in genetic distance with geographic distance locally (Figure 2D, compare to 2F). However, at larger scales (e.g. comparing island to mainland) islands are more dissimilar, as seen by the greater genetic distances at larger geographic distances (Figure 2E, compare to 2G; also see the first two principal components of SNPs, Figure S7). These results highlight the capacity of the method to contrast patterns across scales requiring only dilation of the analyzing kernel.

**Figure 2:**
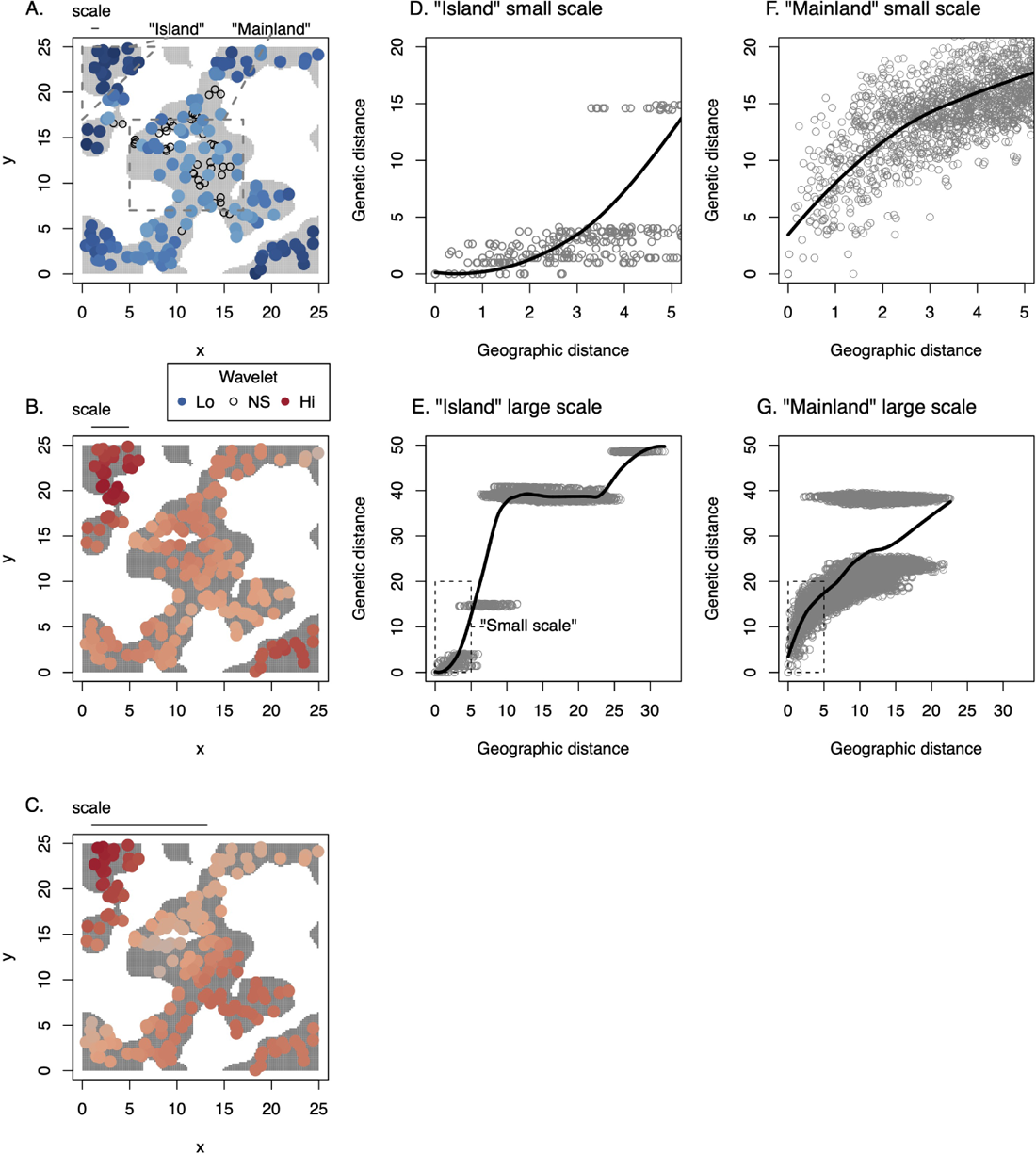
Wavelet genetic dissimilarity identifies scale-specific, localized patterns in a heterogeneous landscape, with pairwise distance plots for comparison. (A-C) Maps of simulated landscape where habitat is gray (in background) and unsuitable areas are white. Sampled individuals are circles. Colors represent sampling locations where wavelet genetic dissimilarity was significantly high (red) or low (blue), with *s*, the wavelet scale, shown at top of each panel as a horizontal line. At the smallest scales (A), samples have less dissimilarity than expected, especially in the island in the upper left of the landscape. This pattern can also be seen (D,F) when comparing pairwise geographic versus euclidean genetic distances for samples in the different regions of the landscape (dashed grey lines in A). At larger spatial scales (B-C), all locations have significantly greater dissimilarity than expected due to limited gene flow. However, the same islands show the greatest dissimilarity at large scales (lower panels), due to their high genetic difference from mainland samples at center. This pattern can also be seen in the pairwise genetic distances across larger geographic distances (E,G). (D-G) Loess smoothing curves are shown.

#### 2.1.5 Simulated neutral patterns in a colonizing and rangeexpanding species

For a second scenario where we expected localized, scale-specific heterogeneity, we simulated an invasion/range expansion. Beyond the importance of invasions in applied biology, the changes in spatial genetic patterns over time are of general interest (Castric and Bernatchez 2003; Le Corre et al. 1997; Slatkin 1991, 1993), considering that all species ranges are dynamic and many “native” species still bear clear evidence of expansion, e.g. following the last glacial maximum.

We simulated invasion across a square landscape of the same size as before, but beginning with identical individuals only in the middle at the bottom edge of the landscape (Figure 3). We sampled 200 individuals at time steps 100, 250, 500, 1000, 1500, 2000, through the full populating of the landscape around 2500 years and until the 3000th time step.

**Figure 3:**
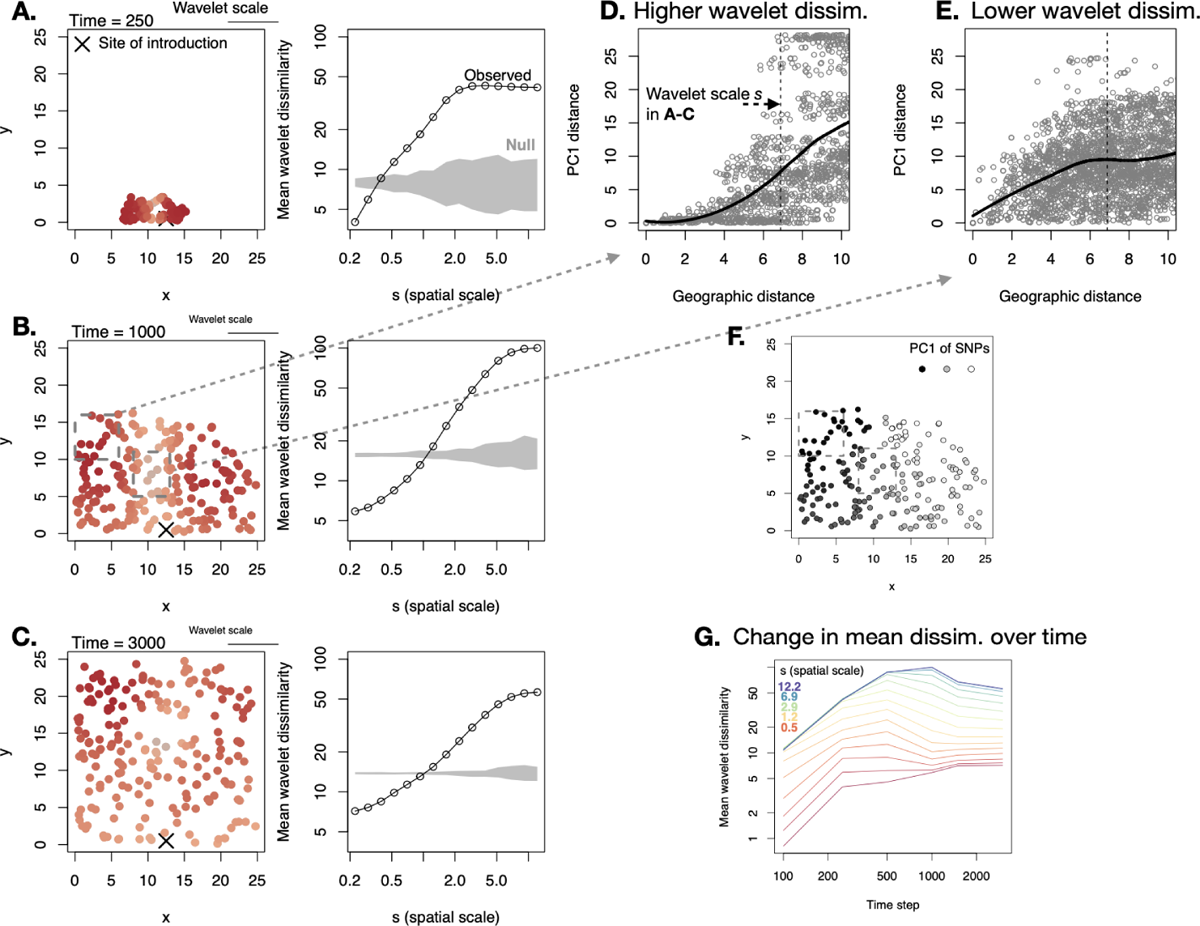
Wavelet genetic dissimilarity reveals dynamic spatial patterns during an invasion across a homogeneous landscape. Left column of panels (A-C) shows a map of the landscape through time, with 200 sampled individuals at each time step and the wavelet dissimilarity at *s* = 6.9 at their location. Darker red indicates greater wavelet dissimilarity. In the second time step, 1000, two regions are highlighted in dashed boxes (B), one with higher dissimilarity at *s* = 6.9 (D) and one with lower dissimilarity at this scale (E). (D-E) show pairwise geographic distance versus distance in the first PC of SNPs for samples from these regions. (F) shows the loadings of each sample on the first PC of SNPs. (D-E) highlight the greater increase in PC1 distance with geographic distance at this scale (vertical dashed lines) in (D), compared to the smaller increase in PC1 distance at this scale in (E). In particular, the region highlighted in (D) is homogeneous at short distances but very distinct at distances at the highlighted scale *s* = 6.9, indicating the major genetic turnover at this scale and location. (G) Mean wavelet dissimilarity across the landscape changes over time, highlighting the dynamic spatial population genetic patterns across invasions. Loess smoothing curves are shown in (E-F).

We characterized wavelet genetic dissimilarity and found substantial heterogeneity across different regions and across time (e.g. for *s* = 6.9, dark versus light red in Figure 3A-C). This heterogeneity in genetic turnover can be seen by contrasting genotypes from different regions. Near the expansion front, there is relative homogeneity and low diversity locally in new populations, but with rapid turnover in genotypes separated by space, resulting in high wavelet dissimilarity at intermediate spatial scales (Figure 3D). In the range interior, there is greater local diversity and less turnover in genotype across space, i.e. a weaker isolation by distance (Figure 3E, see all SNP genetic distance plot Figure S8). Supporting the role of founder effects and low diversity at expanding range margins in driving these patterns, we observed a decline in mediumand large-scale wavelet dissimilarity in later years (Figure 3G) after the landscape had been populated.

These patterns highlight how wavelet dissimilarity is capturing scale-specific turnover in genetic composition, rather than merely genetic distance at a given geographic distance. Comparing the two regions highlighted in Figure 3B, the genetic distances at a geographic distance of 6.9 are not strikingly different (Figure S8). Rather what distinguishes these regions is their rate of genetic change in composition at this scale, as highlighted in Figure 3. The region of high wavelet dissimilarity at *s* = 6.9 (Figure 3B) transitions from homogeneity among nearby samples to high genetic distance at larger scales (Figure 3D, S8). By contrast the region of low wavelet dissimilarity at *s* = 6.9 (Figure 3B) starts out with greater genetic distance among nearby samples with a modest increase in genetic distance at larger scales (Figure 3E, S8).

Overall, these simulations show the capacity of *D^wav^*(*s*), wavelet genetic dissimilarity, to capture localized, scale specific trends in genetic composition. Given the spatial heterogeneity in nature and the dynamics of populations and species ranges through time, there are likely many such patterns waiting to be described to shed light on patterns of gene flow and population history.

### 2.2 Finding the loci of local adaptation

#### 2.2.1 Using wavelet transforms to identify outliers of spatial pattern in allele frequency

We can also use our approach to transforming allele frequencies to identify particular genetic loci involved in local adaptation, and the regions and spatial scales of turnover in their allele frequency. Our strategy is (as before) to first calculate (*T^wav^f_i_*)(*a, b, s*), the wavelet transform, for each locus *i* at each sampling point *a, b* for a set of chosen spatial scales *s ∈ S*.

Because of different ages and histories of drift, mutations will vary in their global allele frequency and thus global variance. To facilitate comparisons among loci for relative evidence of selection, we can normalize spatial patterns in allele frequency by total variation across locations, as is done when calculating *F_ST_*.

Here we divide the wavelet transforms of allele frequency by the standard deviation of global allele frequency variation for each locus *i*, *sd*(*f_i_*). This normalization is greatest when minor allele frequency is 0.5 for a biallelic locus, and yields a scaled wavelet transformed allele frequency: (*T^wav^f_i_*)(*a, b, s*)*/sd*(*f_i_*), for a given location and scale.

We then calculate the variance across sampling locations of (*T^wav^f_i_*)(*a, b, s*)*/sd*(*f_i_*) and refer to this quantity as the “scale-specific genetic variance.” This scaled-specific variance is akin to *F_ST_* in being a measure of spatial variation in allele frequency normalized to total variation (which is determined by mean allele frequency). High scale-specific variance for a given locus indicates high variation at that scale relative to the total variation and mean allele frequency. We then used a *χ*^2^ null distribution across all genomic loci to calculate parametric p-values (Cavalli-Sforza 1966; Lewontin and Krakauer 1973) and used the approach of Whitlock and Lotterhos (2015) to fit the degrees of freedom of the distribution of scale-specific genetic variances (see Supplemental Methods). Applying this approach to a range of simulated scenarios as well as an empirical dataset (described below), we see that the *χ*^2^ distribution with a maximum likelihood fit to determine degrees of freedom provides a reasonably close fit to the distribution of scale-specific genetic variance among SNPs (Figures S9-S12).

#### 2.2.2 Simulated local adaptation

First, we present some specific individual simulations for illustration, and then a larger set with more variation in underlying parameters. We simulated a species with the same life history parameters as in simulations above, with the addition of spatially varying viability selection on a quantitative trait. We imposed two geometries of spatially varying selection, one a linear gradient and the other a square patch of different habitat selecting for a different trait value. As with the neutral simulations, simulations with selection began with organisms distributed across the landscape, with an ancestral trait value of zero. In these simulations, 1% of mutations influenced the quantitative trait with additive effects and with effect size normally distributed with a standard deviation of 5. For the linear gradient, the optimal trait value was 0.5 at one extreme and −0.5 at the other extreme, on a 25×25 square landscape. Selection was imposed using a Gaussian fitness function to proportionally reduce survival probability, with standard deviation *σ_k_*. In this first simulation, *σ_k_* = 0.5. Carrying capacity was roughly 5 individuals per square unit area, and simulated populations usually stabilized close to this density. Full details of simulation, including complete code, can be found in supplemental materials and on GitHub (https://github.com/jesserlasky/WaveletSpatialGenetic).

In the first simulation along a linear gradient, there were 2 selected loci with minor allele frequency (MAF) at least 0.1, where the scale of mating and propagule dispersal were each *σ* = 1.1, after 2,000 time steps. The two loci under stronger selection were clearly identified by the scale-specific genetic variance *var*((*T^wav^f_i_*)(*a, b, s*)*/sd*(*f_i_*)) at the larger spatial scales (Figure 4). When there is a linear selective gradient across the entire landscape, the largest spatial scale is the one most strongly differentiating environments and the strongest scalespecific genetic variance was at the largest scale (Figure 4). However, power may not be greatest at these largest scales, because population structure also is greatest at these largest scales. Instead, power was greatest at intermediate scales, as seen by the lowest p-values being detected at these intermediate scales (Figure 4). At these scales there is greater gene flow but still some degree of changing selection that may maximize power to detect selection.

**Figure 4:**
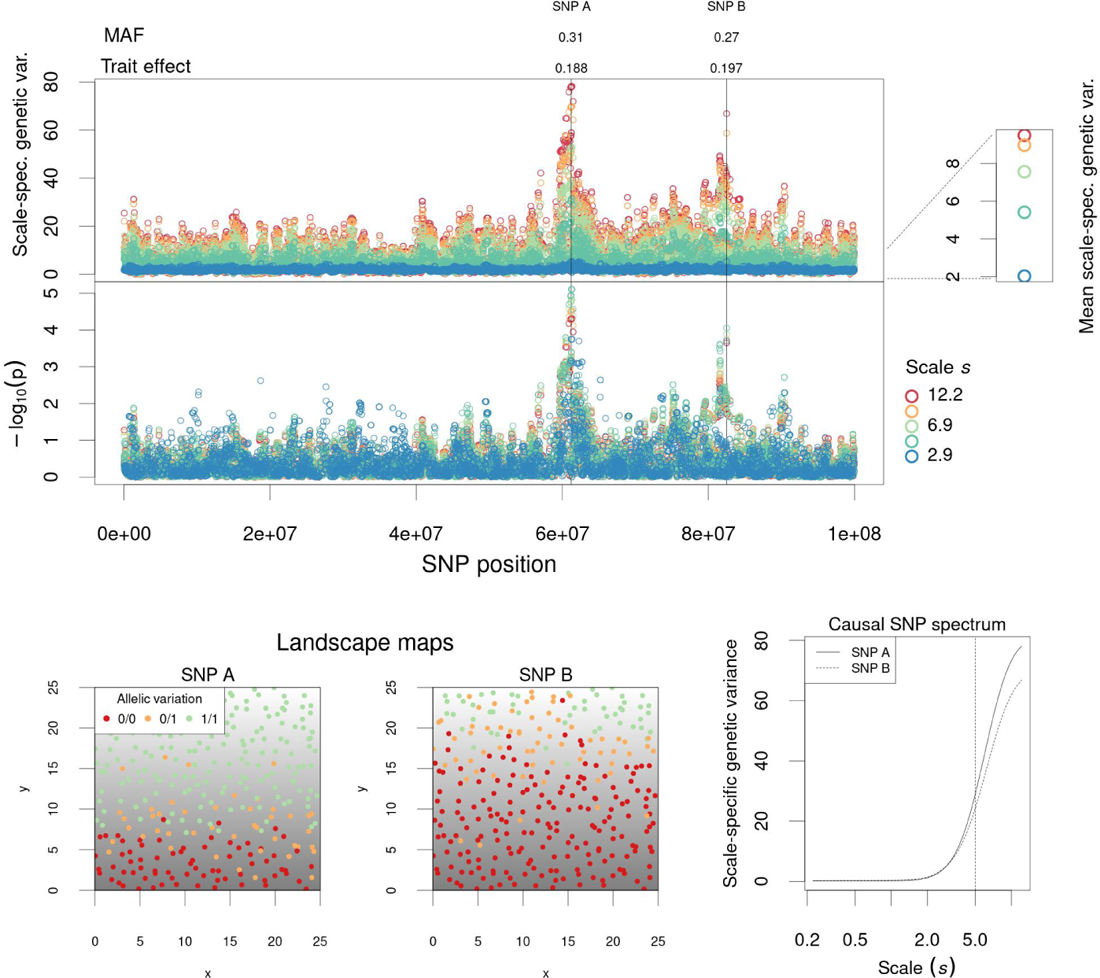
Scale-specific genetic variance test applied to simulations with a linear selective gradient. (top panels) Genome-wide variation in scale-specific genetic variance, *var*((*T^wav^f_i_*)(*a, b, s*)*/sd*(*f_i_*)), for five different scales *s* and upper-tail p-values for *χ*^2^ test using fitted values of d.f. Each point represents a SNP at a specific scale. Loci under selection are indicated with vertical lines along with the absolute value of the derived allele’s effect on the trait and MAF. At bottom are shown maps of the two selected loci as well as their spectra of scale-specific genetic variance. At upper right the mean scalespecific genetic variance across all genomic loci is shown for each scale *s*. The scale of mating and propagule dispersal were each *σ* = 1.1. Gaussian viability selection was imposed with *σ_k_* = 0.5. Carrying capacity was approximately 5 individuals per square unit area.

We next simulated change in selection in a discrete habitat patch, which may more closely correspond to the setting where researchers would find useful a flexible approach to finding spatial patterns in allele frequency, especially if the patches of distinct environment are not known by researchers. In our simulation there was a large central patch, 10×10, that selected for distinct trait values (trait optimum = 0.5) compared to the outer parts of the landscape (trait optimum = −0.5). Selection was initially weakly stabilizing (*σ_k_* = 3 around the optimum of zero for the first 500 years to accumulate diversity, and then the patch selective differences were imposed with stronger selection, *σ_k_* = 0.08. The scales of mating and propagule dispersal were each *σ* = 2. Carrying capacity was was roughly 50 individuals per square unit area.

In this simulation we present results after 3000 time steps, where there was a single common QTL under selection (Figure 5). We found several spurious large scale peaks in scale-specific genetic variance (Figure 5A), but when using the *χ*^2^ test on these statistics we clearly identified the single QTL under selection, with lowest p-values for intermediate scales (Figure 5B).

**Figure 5:**
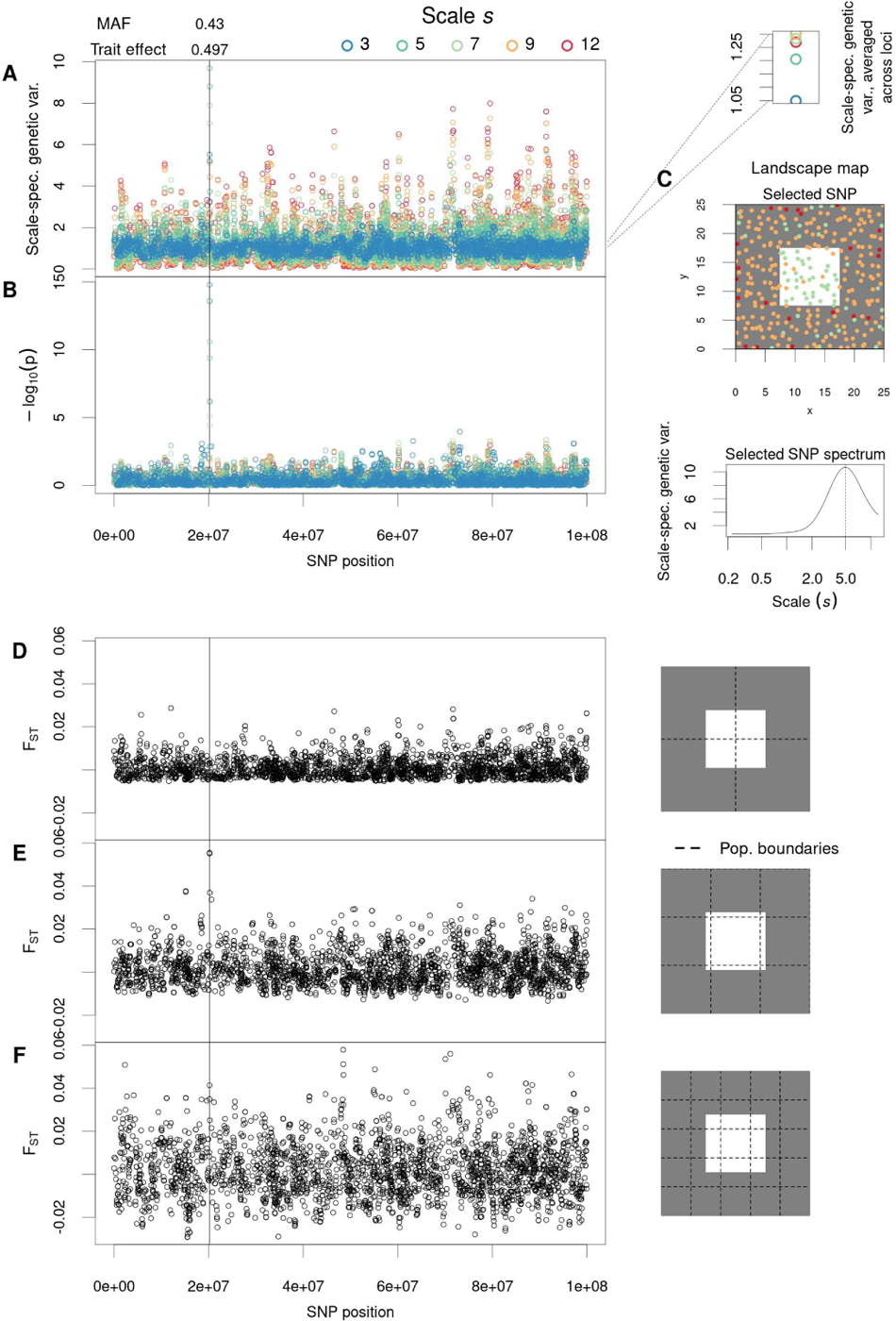
Simulations of local adaptation to a single discrete patch of different habitat. (A) Genome-wide variation in scale-specific genetic variance *var*((*T^wav^f_i_*)(*a, b, s*)*/sd*(*f_i_*)) and (B) *χ*^2^ p-values for six different scales *s*, for a discrete habitat difference after 3000 simulated years. Each point in the left panels represents a SNP, and wavelet statistics (A-B) at specific scales. The selected SNP is indicated with a vertical line along with the absolute value of a derived allele’s effect on the trait and MAF. (C) A map of the landscape with individuals’ genotypes at the causal SNP indicated with color, in addition to the spectrum of scale-specific genetic variance at this SNP, showing a peak at approximately half the patch width (vertical line at 5).(D-E) Implementation of *F_ST_* using arbitrary boundaries for populations. This approach can easily miss causal loci (C,E) if the delineated population boundaries do not match habitat boundaries. (A) At upper right the mean scale-specific genetic variance across all loci is shown for each scale *s*. The1s8cale of mating and propagule dispersal were each *σ* = 2. Gaussian viability selection was imposed with *σ_k_* = 0.08.

We calculated the scale-specific genetic variance across a denser spectrum of scales *s* for the causal SNP, to determine at what scale variance was greatest. We found the maximum scale-specific genetic variance for the causal SNP was at 5.02, approximately half the length of a patch edge (Figure 5C). For illustration, we also calculated *F_ST_* (Goudet 2005; Weir and Cockerham 1984) for several naively discretized subpopulation scenarios for a simple illustration of how results are sensitive to discretization (Figure 5D-F). We also implement our test on these two simulated landscapes but with biased sampling (as described for neutral landscapes above) and found our ability to detect causal loci was robust (Figure S13).

#### 2.2.3 Evaluating the scale-specific genetic variance test

As an initial assessment of the general appropriateness of the scale-specific genetic variance test we proposed above, we conducted additional simulations on two types of landscapes with varying life history parameters. These simulations were not meant to be an exhaustive evaluation of the performance of this new test; we leave a more extensive evaluation for future studies.

Here, we again used the discrete habitat patch landscape and the linear gradient landscape but with a wider range of parameter variation. We tested a range of mating and dispersal (*σ*) scales including 0.25, 0.5, 1, and 2, and a range of stabilizing selection (*σ_k_*) values including 0.125, 0.25, 0.5, and 1. Three simulations were conducted for each combination of parameter settings and each ran for 10,000 years.

Because PCAdapt is one of the few methods for identification of spatial pattern in allele frequency that does not require subpopulation discretization and in theory could detect patterns at multiple scales, we also implemented this method. We used the PCA of the scaled genotype matrix, thinned for LD but including causal SNPs, to extract the z-scores and p-values of each SNP with a cutoff of p = 0.05. We used a scree plot showing the percentage of variance explained in decreasing order to identify the optimal number of principal components following Cattell’s rule (Duforet-Frebourg et al. 2016).

Calculating false and true positive rates for PCAdapt was straightforward, but for the scale specific genetic variance test there are several tests (one at each scale) for each SNP. To conservatively represent inference across these multiple tests, we considered SNPs a significant result if one of the tested scales was significant. Because the individual scale tests are slightly conservative, and continuous wavelet transforms are correlated across scales (and hence not completely independent tests), we expected the resulting false positive rates would not be unduly high. Overall the scale-specific genetic variance test showed good false positive rates. Across simulations, the proportion of SNPs with *χ*^2^ upper-tail *p <* 0.05 at one scale was usually close to but sometimes slightly more than 0.05 (Figure 6). By contrast, under scenarios of low gene flow and strong stabilizing selection, nominal false positive rates were high for PCAdapt, often *>* 0.15.

**Figure 6:**
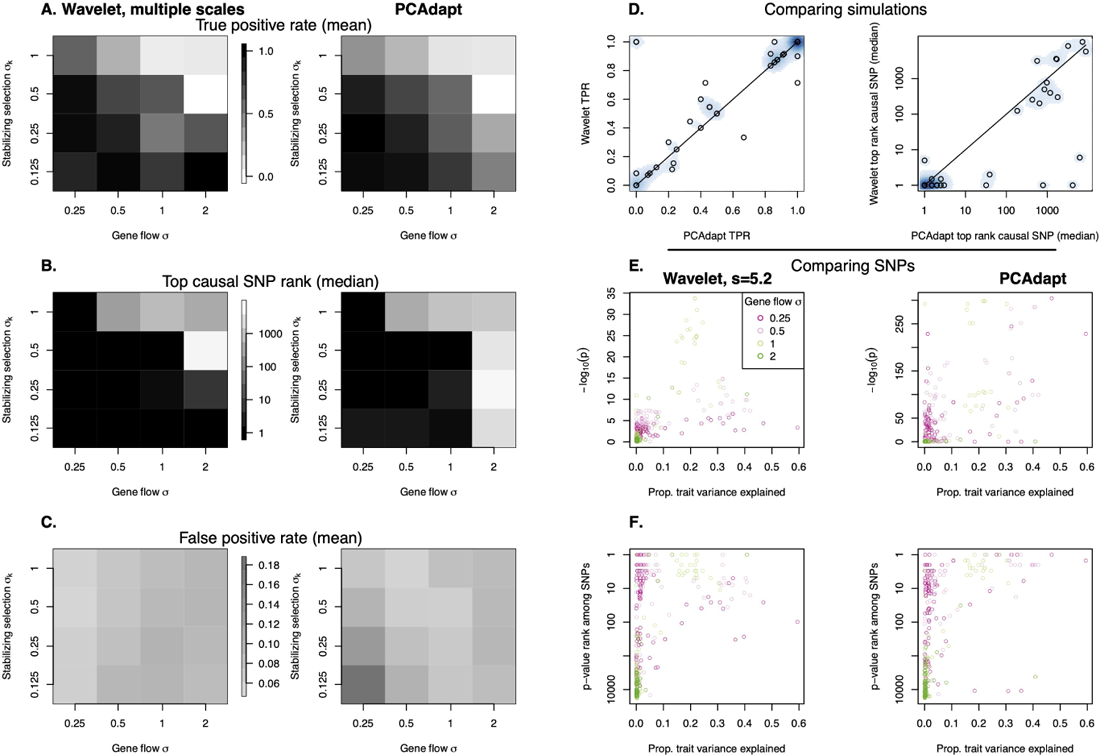
Comparing the scale-specific genetic variance test with PCAdapt in simulations of adaptation to single discrete patch of different habitat. (A) True positive rates (nominal *p <* 0.05) for each combination of simulation parameters, the scales of mating and dispersal *σ* and the standard deviation of the Gaussian stabilizing selection function *σ_k_*. (B) An alternate view of statistical power based on the median rank of the top selected SNP among all SNPs. (C) False positive rates (nominal *p <* 0.05). (D) Comparing power between the two statistical approaches for the different simulation runs. Density of points is shown in the blue scale so as to indicate where many simulations had the same result. The line indiciates a 1:1 relationship. (E-F) Individual selected SNPs in simulations, showing their nominal *p* values and ranks among all SNPs, colored based on *σ* in the simulation. The x-axis represents the proportion of total phenotypic variation among sampled individuals that was explained by the given SNP (*R*^2^ from a linear model).

Power to detect SNPs (proportion of selected SNPs with *p <* 0.05) under selection was generally high (true positive rate near 1) but sometimes low, depending on the strength of selection (*σ_k_*) and mating and dispersal scales (*σ*) (Figure 6). When gene flow was high and selection was weak, power was low for both the scale-specific genetic variance test and PCAdapt. This also corresponds to the scenario when local adaptation is weakest (Kirkpatrick and N. H. Barton 1997). In addition to considering power simply based on *p* for each SNP, we also considered power using the top *p*-value rank among selected SNPs under each simulation, based on the reasoning that researchers may want to follow up on top ranked outlier SNPs first before any lower ranked SNPs. This approach showed similar results, with high power for both the scale-specific genetic variance test and PCAdapt except when gene flow was high and selection weak. In general, the two methods showed comparable power across different scenarios (Figure 6), with some indication that the scale-specific genetic variance test had higher power under high gene flow and PCAdapt slightly higher power under lower gene flow. By plotting individual SNPs we can see that for the upper end of gene flow scenarios (*σ* = 1 or 2), the scale-specific genetic variance test more consistently identified selected SNPs at the top compared to PCAdapt. For the low gene flow scenarios, PCAdapt more consistently identified large effect variants, while the scale-specific genetic variance test more consistently identified the smaller effect variants (see results for linear gradient in Figure S14). Overall, the similarities in true and false positive rates between methods suggest that our wavelet approach is effective compared to other related tools, while our test also offers the ability to explicitly consider variation in spatial scale.

### 2.3 Testing for spatial pattern in quantitative trait loci (QTL)

When testing for spatially-varying selection on a quantitative trait, one approach is ask whether QTL identified from association or linkage mapping studies show greater allele frequency differences among populations than expected (Berg and G. Coop 2014; Price et al. 2018). Here we implement such an approach to compare wavelet transformed allele frequencies for QTL *L* to a set of randomly selected loci of the same number and distribution.

For this test we calculate the mean of scale-specific genetic variance for all QTL with MAF at least 0.05 among sampled individuals. We then permute the identity of causal QTL across the genome and recalculate the mean scalespecific genetic variance, and repeat this process 1000 times to generate a null distribution of mean scale-specific genetic variance of QTL for each scale *s*.

We illustrate this test here briefly using a simulation of adaptation to a square patch of habitat in the middle of a landscape, with the two gene flow parameters *σ* = 0.5, the strength of selection *σ_K_* = 0.5, carrying capacity *∼* 5 individuals per square unit area. After 1000 generations we sampled 300 individuals, from which there were 13 QTL for the trait under selection with MAF at least 0.05. We then calculated the mean scale-specific genetic variance for these QTL across scales *s* and compared to the null permutations of randomly selected 13 SNPs from the genome.

We found significantly higher mean scale-specific genetic variance for the QTL than the null expectation at all 6 scales tested. Although the scale-specific genetic variance was greatest at the largest scales for the QTL, these scales did not show as great a distinction when comparing to the null. The greatest mean wavelet variance of QTL relative to null came at the intermediate scales of 3-5, which was approximately 1/3-1/2 the width of the habitat patch (Figure S15).

### 2.4 Application to an empirical system

#### 2.4.1 Genome-wide wavelet dissimilarity

We applied our approach to an empirical dataset of diverse, broadly distributed genotypes with whole genome resequencing data: 908 genotypes from 680 natural populations of the model plant, *Arabidopsis thaliana* (Brassicaceae). We used a published Arabidopsis dataset (Alonso-Blanco et al. 2016), only including Eurasian populations and excluding highly distinct “relicts” and also likely contaminant accessions (Pisupati et al. 2017). For locations with more than one accession genotyped we calculated allele frequency. We used a total of 129536 SNPs filtered for minor allele frequency (MAF*>* 0.05) and LD (Zheng et al. 2012).

We first calculated the genome-wide wavelet dissimilarity, *D^wav^*(*s*), across a series of increasing scales *s* at even intervals in log distance units from *∼* 50 m to approximately half the distance separating the farthest samples, *∼* 3000 km. We observed increasing mean genome-wide wavelet dissimilarity at larger scales (Figure 7), a pattern indicative of isolation by distance, on average, across the landscape. Arabidopsis showed significantly low dissimilarity at scales less than *∼* 5 km, likely due to the homogenizing effect of gene flow. However, we found significantly high dissimilarity at scales greater than *∼* 7 km. This scale of significantly high dissimilarity may be a relatively short distance, considering that Arabidopsis is largely self pollinating and lacks clear seed dispersal mechanisms (though seeds of some genotypes form mucus in water that increases buoyancy) (Saez-Aguayo et al. 2014). At scales greater than *∼* 120 km we found an increase in the slope relating scale *s* and dissimilarity, perhaps signifying a scale at which local adaptation begins to emerge.

**Figure 7:**
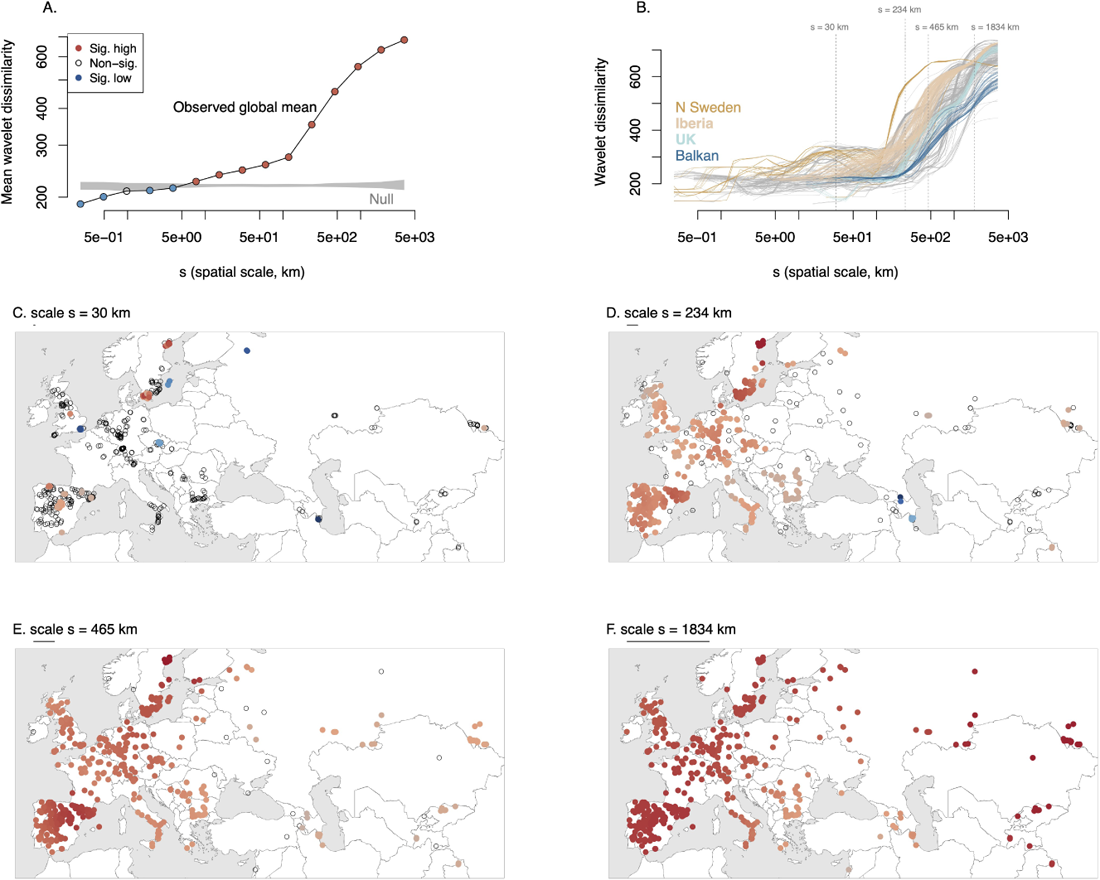
Genome-wide wavelet dissimilarity, *D^wav^*(*s*)**, for Arabidopsis genotypes.** (A) The global mean dissimilarity across scales compared to the null expectation (gray ribbon) and (B) the dissimilarity across scales centered on each sampled genotype, with several regions highlighted (vertical lines indicate scales shown in panels C-F). (C-F) Selected scales highlight the changes in dissimilarity across locations, with each circle indicating a genotyped sample/population. Red indicates significantly greater wavelet dissimilarity than expected, blue significantly less than expected. For the map panels, the intensity of color shading indicates the relative variation (for a given scale) in *D^wav^*(*s*) among significant locations.

The locations of scale-specific dissimilarity among Arabidopsis populations revealed several interesting patterns. Even by the *∼* 30 km scale, there were three notable regions of significantly high dissimilarity: northern Spain and extreme southern and northern Sweden (Figure 7). The high dissimilarity at this scale in northern Spain corresponds to the most mountainous regions of Iberia, suggesting that limitations to gene flow across this rugged landscape have led to especially strong isolation among populations at short distances. In northern

Sweden, Long et al. (2013) previously found a particularly steep increase in isolation-by-distance. Alonso-Blanco et al. (2016) found that genetic distance was greatest among accessions from Southern Sweden at scales from *∼* 20 *–* 250 compared to regions farther south. At larger, among-region scales, dissimilarity was significantly high across the range, with Iberia and northern Sweden again being most dissimilar at *∼* 234 km and surpassed by central Asia at *∼* 1834 km as being most dissimilar. Iberia and northern Sweden contain many accessions distantly related to other accessions, likely due to isolation during glaciation and subsequent demographic histories (Alonso-Blanco et al. 2016). This scale in Asia separates populations in Siberia from those further south in the Tian Shan and Himalayas, indicating substantial divergence potentially due to limited gene flow across the heterogeneous landscape. By contrast, populations in the UK and the Balkan peninsula had low dissimilarity across a range of scales, possibly due to reduced diversity and a more recent history of spread in these regions.

#### 2.4.2 Identifying putative locally-adapted loci

For this analysis, we used the same genotypes as in the prior section but not filtered for LD, leaving 1,642,040 SNPs with MAF*>* 0.1 (Alonso-Blanco et al. 2016). The scale-specific genetic variance test identified putative locally adapted loci (Figure S16). The distribution of scale-specific genetic variance among SNPs was reasonably matched to the theoretical *χ*^2^ distribution (Figure S12).

Among notable loci, at the *∼* 59 km scale, the #2 QTl and #3 SNP is in the coding region of METACASPASE 4 (MC4), a gene that controls biotic and abiotic stress-induced programmed cell death (Hander et al. 2019; Shen, Liu, and Li 2019). To speculate, if MC4 were involved in coevolution with microbial pathogens we might expect rapid allele frequency dynamics and thus a pattern of high variation among even nearby populations.

The #1 SNP for the *∼* 282 km scale was in the coding sequence of the DOG1 gene (Figure 8, Figure S12). This SNP, Chr. 5, 18,590,741 was also strongly associated with flowering time (see next section) and germination and tags known functional polymorphisms at this gene that are likely locally adaptive (Martínez-Berdeja et al. 2020). The spatial pattern of variation at this locus (Figure 8) is complicated, highlighting the benefit of the flexible wavelet approach. By contrast, imposing a grid on this landscape, or using national political boundaries to calculate *F_ST_* could easily miss the signal as did Horton et al. (2012). The climate-allele frequency associations for DOG1 are also complicated and non-monotonic (**gamba•genomics•2023**; Martínez-Berdeja et al. 2020), making it challenging for genotype-environment association approaches (Jesse R Lasky, Emily B Josephs, and Morris 2023).

**Figure 8:**
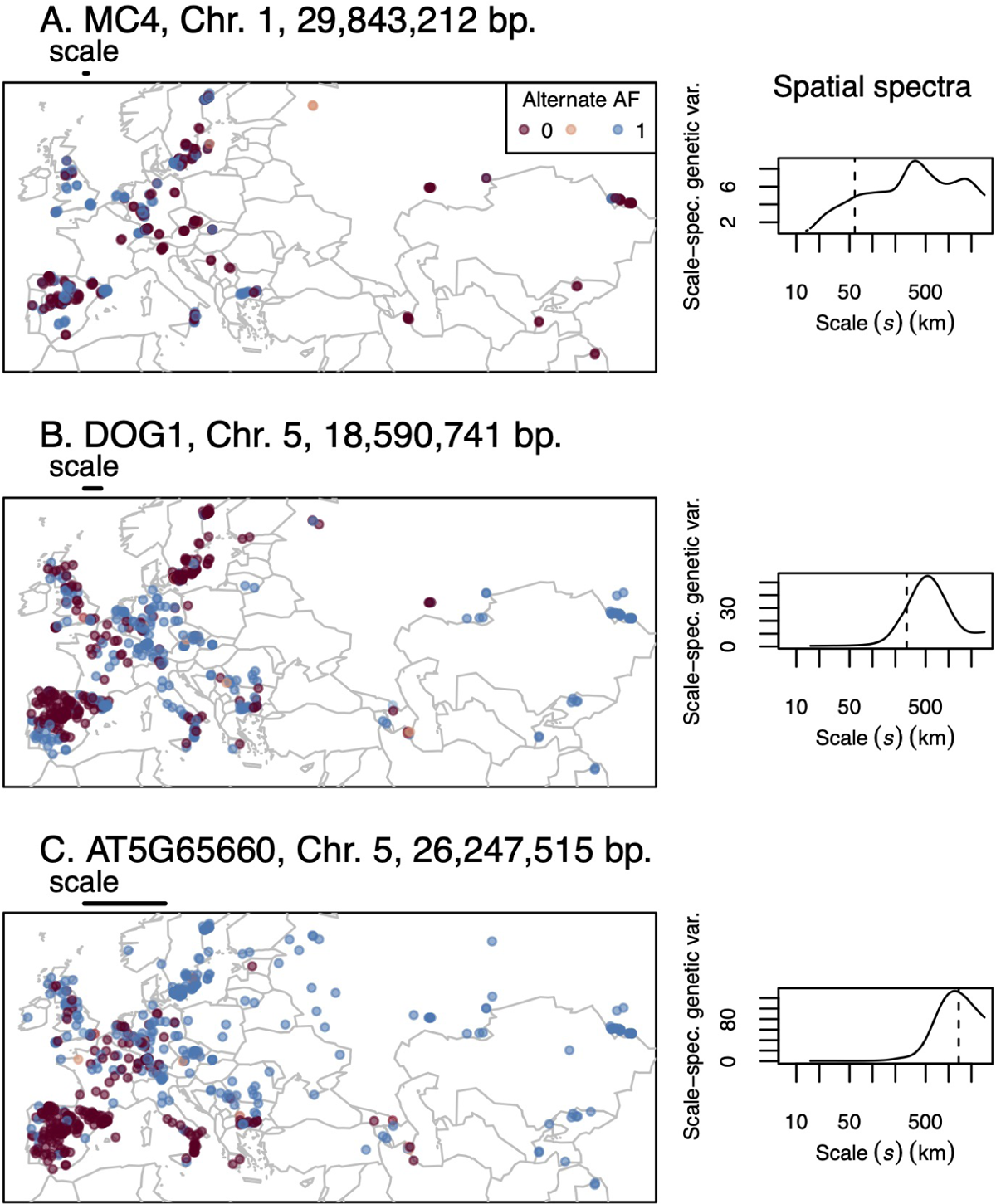
Allelic variation (colors) for SNPs that were top outliers for scale-specific genetic variance test at different scales. On maps at left, the scale for which a SNP was an outlier is indicated by a bar above each map. The right panels show the spatial spectra for each SNP, i.e. the scale-specific genetic variance across a range of scales. Dashed lines indicate the scale for which a SNP was an outlier.

At the *∼* 1359 km scale, the #1 SNP (and also the lowest p-value SNP among all scales, Figure 8, Figure S16) was on chromosome 5 at 26,247,515 bp, 555 bp upstream from AT5G65660, a hydroxyproline-rich glycoprotein family protein. These are cell wall glycoproteins with important roles in development and growth (Johnson et al. 2017) some of which have a role in abiotic stress response (Tseng et al. 2013).

#### 2.4.3 Testing for local adaptation in quantitative trait loci (QTL)

We tested for non-random scale-specific genetic variance of QTL for Arabidopsis flowering time, a trait that is likely involved in local adaptation (Ågren et al. 2017). We used previously published data on flowering time: days to flower at 10°C measured on 1003 genotypes and days to flower at 16°C measured on 970 resequenced genotypes (Alonso-Blanco et al. 2016). We then performed mixed-model genome wide association studies (GWAS) in GEMMA (v 0.98.3) (Zhou and Stephens 2012) with 2,048,993 M SNPs filtered for minor allele frequency (MAF*>* 0.05), while controlling for genome-wide similarity among ecotypes.

We found that top flowering time GWAS SNPs showed significantly elevated scale-specific genetic variance at several intermediate spatial scales tested. For flowering time at both 10° and 16°C, scale-specific genetic variance was significantly elevated for the top 1,000 SNPs at the 282, 619, and 1359 km scales, but not always at the largest or smallest scales (Figure 9). In particular the scale-specific genetic variances were greatest for the *∼* 282 km scale where the mean scale specific genetic variance for 16°C QTL was 15.2 standard deviations above the null mean, and the *∼* 619 km scale, where the mean scale specific genetic variance for 10°C QTL was 13.5 standard deviations above the null mean. For QTL from both temperature experiments, results were nearly equivalent if we instead used the top 100 SNPs.

**Figure 9:**
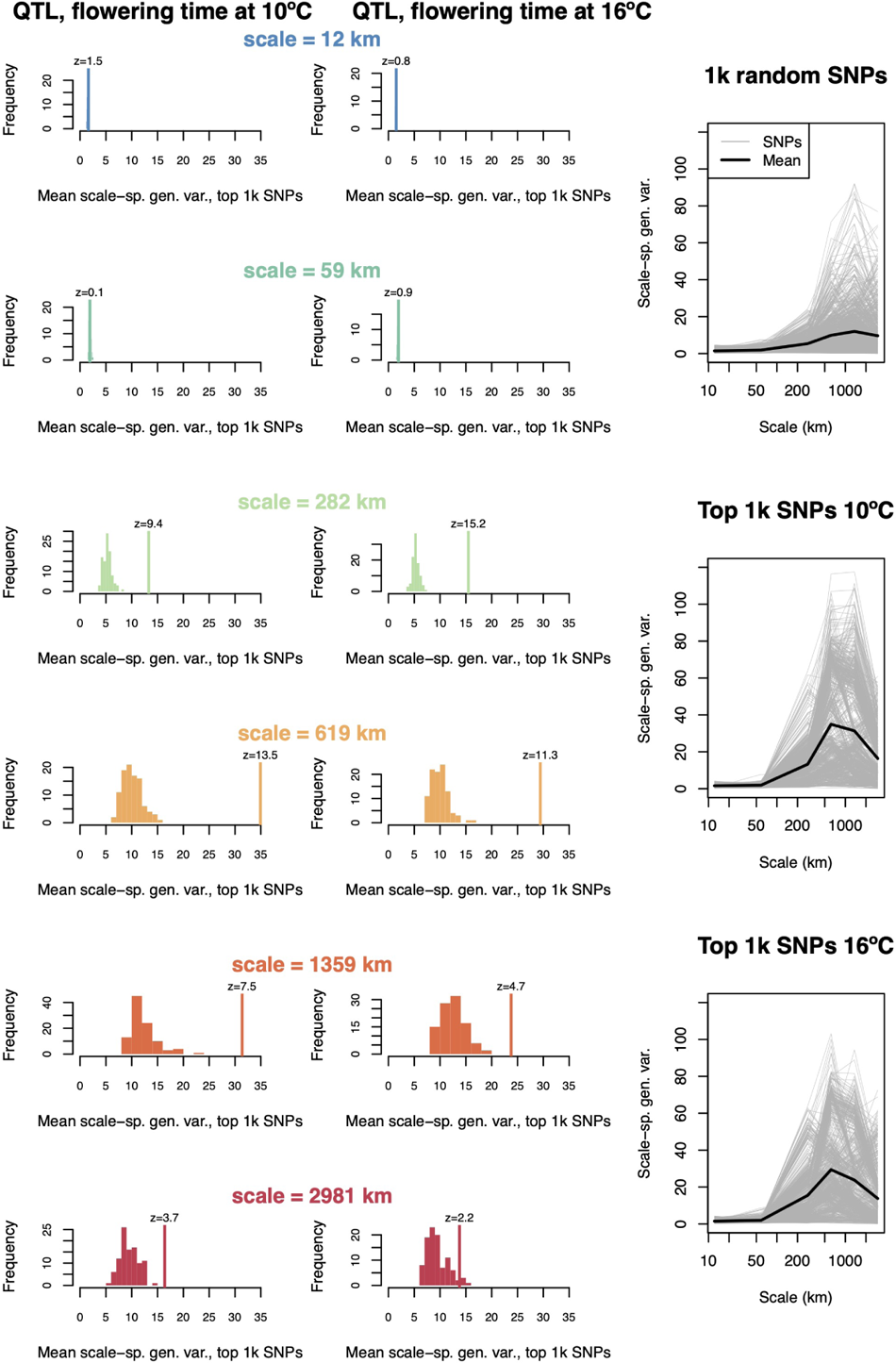
Testing for selection on Arabidopsis flowering time QTL. We compared scale-specific genetic variance, *var*((*T^wav^f_i_*)(*a, b, s*)*/sd*(*f_i_*)), of QTL with random SNPs, for five different scales *s*, for flowering time measured at 10°C and 16°C. The first two columns show the observed mean of the top 1,000 flowering time SNPs with a vertical line and a z-score. The histograms show null distributions of scale-specific genetic variance based on permutations of an equal number of markers with an equal distribution as the flowering time QTL. At right the scale-specific genetic variance is shown for random SNPs and for the flowering time QTL (gray lines), across scales, with the mean indicated by a black line.

## 3 Discussion

Geneticists have long developed theory for spatial patterns in allele frequency (Haldane 1948; Malecot 1948; Wright 1943). Empiricist have sought to use these patterns make inference about underlying processes of demography, gene flow, and selection (Lewontin and Krakauer 1973; McRae et al. 2008; Rousset 2000). While statistical approaches have been developed to characterize geographic patterns, few are flexible enough to incorporate patterns at a range of scales that are also localized in space. Because wavelet transforms have these properties, we think they may be useful tools for geneticists. Here we demonstrated several applications of wavelet transforms to capture patterns in whole genome variation and at particular loci, under a range of neutral and non-neutral scenarios.

Some important existing approaches are based on discretization of spatially-distributed samples into spatial bins, i.e. putative populations (Bishop, Chambers, and I. J. Wang 2023; Petkova, Novembre, and Stephens 2016; Weir and Cockerham 1984). However, without prior knowledge of selective gradients, patterns of gene flow, or relevant barriers, it is often unclear how to delineate these populations. For example, we can see how the specific discretization can hinder our ability to find locally-adapted loci in our simulations (Figure 5) and in empirical studies of Arabidopsis in the case of the phenology gene DOG1 that was missed in previous *F_ST_* scans (Alonso-Blanco et al. 2016; Horton et al. 2012). Our goal in this paper was to provide a new perspective on spatial population genetics using the population-agnostic, and spatially smooth approach of wavelet transforms. We showed how these transforms characterize scale-specific and localized population structure across landscapes (Figures 2, 3, 7). We also showed how wavelet transforms can capture scale-specific evidence of selection on individual genetic loci (Figures 4, 5, 6, 8) and on groups of quantitative trait loci (Figure 9). Our simulations and empirical examples showed substantial heterogeneity in the scale and stationarity of spatial patterns. For example, the wavelet genetic dissimilarity allowed us to identify regions near a front of range expansion with steeper isolation by distance at particular scales due to drift (Figure 3). Additionally, we identified loci underlying local adaptation and showed an example where the evidence for this adaptation was specific to intermediate spatial scales (Figure 5). While existing approaches to characterizing population structure or local adaptation have some ability to characterize scale specific patterns, e.g. those based on ordinations of geography (Wagner, Chávez-Pesqueira, and Forester 2017) or SNPs (Emily B. Josephs et al. 2019), and some can capture localized patterns (e.g Petkova, Novembre, and Stephens 2016), there are few examples of approaches that merge both abilities (Wagner, Chávez-Pesqueira, and Forester 2017).

Like many methods in population genetics that rely on inference from observational data, we view our approaches as exploratory and hypothesis generating. Heterogeneous patterns of genome-wide wavelet dissimilarity suggest demographic hypotheses, some of which can be tested with detailed ecological and genetic study (e.g. Keeley et al. 2017). For genome-scans for loci involved in local adaptation, the p-values resulting from multiple tested scales are comparable and so we recommend starting with the loci having the lowest p-value, and using these to develop hypotheses for functional follow up experiments (Jesse R Lasky, Emily B Josephs, and Morris 2023).

The test for spatial pattern in individual loci we developed owes greatly to previous work from Lewontin and Krakauer (1973) who initially developed *χ*^2^ tests applied to the distribution of *F_ST_* values, and from Whitlock and Lotterhos (2015)’s approach of inferring the degrees of freedom of the *χ*^2^ distribution using maximum likelihood and *F_ST_* across loci. The *χ*^2^ distribution underlies a number of related genetic applied across loci (Fraņcois et al. 2016). However, we note that this test may be slightly conservative in some situations (Figure 6). Nevertheless, we believe there were important signs in our work that this *χ*^2^-based scale-specific genetic variance test was valuable. In particular, we found in our simulation of adaptation to a habitat patch that the scale-specific genetic variance was greatest at large spatial scales but at neutral sites, which obscured spatial pattern at the causal locus (Figure 5). When applying the *χ*^2^ test, we were able to clearly map the causal locus while spurious loci with high scale-specific genetic variance fell away because spatial patterns at those loci still fit within the null distribution.

Relatedly, we found in other simulations and our empirical examples that the strongest evidence for local adaptation was often not at the largest spatial scales (Figure 9), even when the selective gradient was linear across the landscape (i.e. the largest scale, Figure 4). This enhanced power at scales sometimes smaller than the true selective gradients may be due to the limited power to resolve true adaptive clines at large scales from the genome-wide signal of isolation by distance at these scales. At intermediate scales, there may be a better balance of sufficient environmental variation to generate spatial pattern versus higher relatedness between locations due to gene flow.

We note that there remain several limitations to our approach proposed here. First, the ability of wavelet transforms to capture patterns depends on the correspondence between the wavelet form (shape) and the form of the empirical patterns we seek to enhance, and there may be better functional forms to filter spatial patterns in allele frequency. Generally speaking, a more compact smoothing kernel with minimum weight in the tails will be better at revealing abrupt spatial transitions, but at the necessary cost of less precise determination of scale (Heisenberg 1927). Smoothing kernels such as the tricube (*k_x_ ≃* [1 *− x*^3^]) have been shown to optimize certain trade-offs in this space and could be used to construct a difference-of-kernels wavelet. However, the overall influence of kernel shape tends to be much less than the influence of kernel bandwidth in our experience. Second, we have not yet implemented localized tests for selection (i.e. specific to certain locations) as we did with genome-wide dissimilarity. A challenge applying this test at individual loci is that there is a very large number of resulting tests from combinations of loci, locations, and scales. Therefore we have not fully exploited the localized information we derive from the wavelet transforms.

There are number of interesting future directions for research on wavelet characterization of spatial pattern in evolutionary biology. First, we could apply the wavelet transforms to genetic variation in quantitative traits measured in common gardens, to develop tests for selection on traits akin to the *Q_ST_ F_ST_* test (Emily B. Josephs et al. 2019; Whitlock and Guillaume 2009). Second, we could follow the example of Al-Asadi et al. (2019) and apply our measures of genetic dissimilarity to haplotypes of different size to estimate relative variation in the age of population structure. Third, we should test the performance of our tools under a wider range of demographic and selective scenarios to get a more nuanced picture of their strengths and weaknesses. Fourth, null models for wavelet dissimilarity could be constructed using knowledge of gene flow processes (instead of random permutation) to identify locations and scales with specific deviations from null patterns of gene flow.

### 3.1 Conclusion

Population genetics (like most fields) has a long history of arbitrary discretization for the purposes of mathematical, computational, and conceptual convenience. However, the real world often exists without clear boundaries between populations and where processes act simultaneously at multiple scales. We believe that wavelet transforms are one of a range of tools that can move population genetics into a richer but still useful characterization of the natural world.

## 4 Materials and methods

### 4.1 Simulations with SLiM

We developed our simulations by building off the spatial neutral simulation model of Battey et al. C. Battey, Peter L Ralph, and Kern 2020 and model recipes in the SLiM software Haller and Messer 2019 for spatially varying selection on quantitative traits. Scripts are freely available at https://github.com/jesserlasky/WaveletSpatialGenetic/. Parameters differed among scenarios as described previously. The genome had 10^8^ positions with a mutation rate of 10*^−^*^7^. For the initial simulations to demonstrate patterns detectable by our test (Figures 4 and 5), the recombination rate was 10*^−^*^8^ per generation. For the simulations used to compare true positive and false positive rates of the scale-specfic genetic variance test and PCAdapt (Figure 6), we increased the recombination rate to 10*^−^*^7^ per generation and thinned SNPs for LD (while retaining selected SNPs) so as to calculate meaningful SNP-level positive rates across the genome. The carrying capacity per unit square area was roughly 25 for many simulations, unless otherwise noted in text (also see code). The fecundity of individuals in each year was a draw from a Poisson distribution with mean 0.25. Competition occurred among neighbors, potentially resulting in an increased probability of mortality above the 5% background probability (C J Battey, Peter L Ralph, and Kern 2020).

### 4.2 Null model test for wavelet transformed patterns at individual loci

Even under neutrality, individual loci differ in their history and thus not all have identical spatial patterns. To develop a null expectation for the distribution of scale-specific genetic variance across loci, we use the basic approach of Cavalli-Sforza (1966) and Lewontin and Krakauer (1973). Lewontin and Krakauer (1973) used *χ*^2^ null-model tests for *F_ST_* values across multiple loci. The distribution of the sum of squares of *n* independent standard normal variables is *χ*^2^ with *n −* 1 degrees of freedom, so that *^F^*^^^*^ST^* ^(^*^n−^*^1)^ is also *χ*^2^ distributed with *n −* 1 degrees of freedom where *n* is the number of populations and *F*^—^*_ST_* is the mean *F_ST_* among loci (Lewontin and Krakauer 1973). However, the assumption of independence among variables (here, allele frequencies among populations) is often violated, and they instead are embedded in different locations in a heterogeneous (but usually unknown) metapopulation network (Lewontin and Krakauer 1975; Nei and Maruyama 1975; Robertson 1975; Whitlock and Lotterhos 2015).

To solve this problem of non-independence among populations we use the same strategy that Whitlock and Lotterhos (2015) applied to *F_ST_*: we use the distribution of scale-specific genetic variances for each locus to infer the effective number of independent populations (giving the degrees of freedom) for the *χ*^2^ distribution. We used the Whitlock and Lotterhos (2015) method: we trimmed outliers (here the bottom 2.5% SNPs for scale-specific genetic variance) in scale-specific genetic variance, for each scale *s*, then used maximum likelihood to infer the number of independent populations (using the *χ*^2^ maximum likelihood estimation of Whitlock and Lotterhos (2015)), recalculated outliers, and then refit the *χ*^2^ distribution iteratively. Mean scale-specific genetic variance was also calculated in this process while excluding SNPs in the bottom 2.5% tail as well as those with significantly high scale-specific genetic variance at FDR = 0.05. We then used that estimate of the number of effective independent populations to determine the null *χ*^2^ distribution for scale-specific genetic variance.

We then used this null distribution to calculate upper tail probabilities as one-sided p-values, and then used Benjamini Hochberg FDR to get q-values. We found (like Whitlock and Lotterhos 2015) that the *χ*^2^ distribution was sensitive to the inclusion of low MAF variants and thus we also excluded any SNPs with MAF *<* 0.1.

### 4.3 Data availability

Code used to generate the simulations and analyses shown here are freely available at https://github.com/jesserlasky/WaveletSpatialGenetic/.

## Acknowledgments

This work was supported by NIH award R35GM138300 to JRL. We thank Emily Josephs and Benjamin Peter for helpful comments.

## 5 Supporting information

**Fig. S1.**
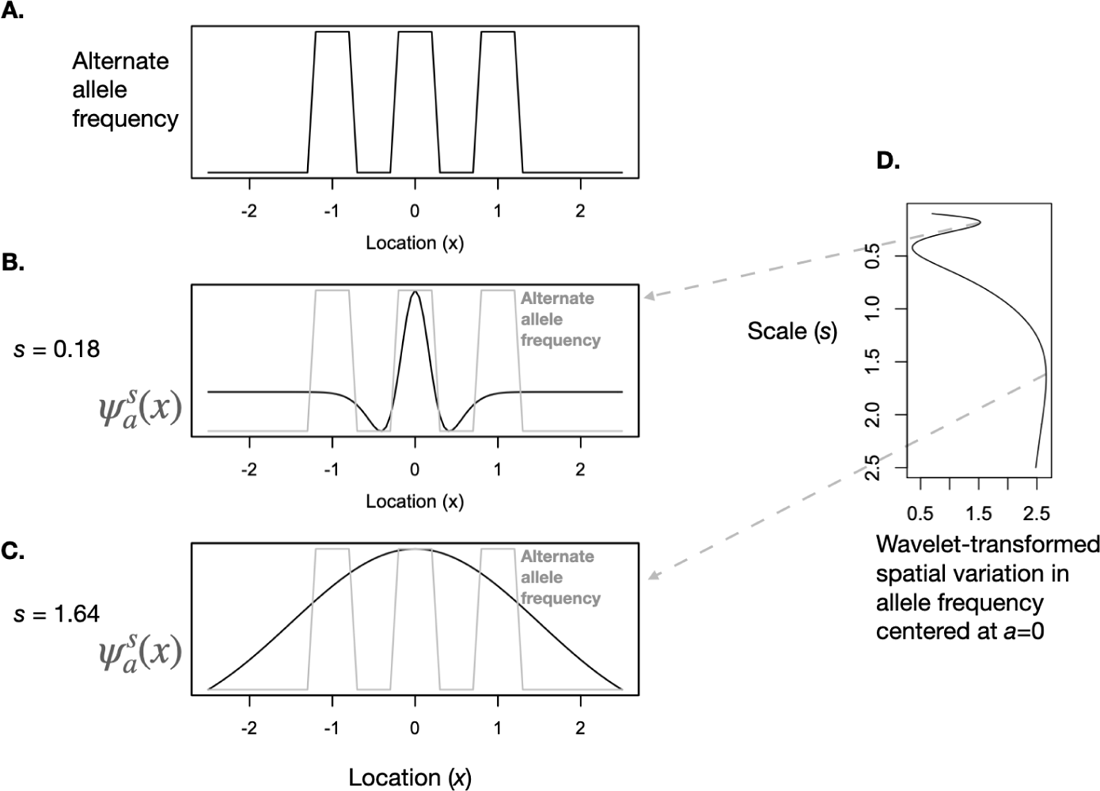
An example of applying a difference of Gaussians (DoG) wavelet to spatial allele frequency patterns (here in one dimension). (A) shows the change in allele frequency across the spatial dimension *x*. (BC) show two DoG wavelets (black curves) of two different scales *s*, centered at location *a* = 0, with the allele frequency pattern overlain in gray. The two selected scales (B-C) are shown because they are the scales at which variation across space in the wavelet transformed allele frequency, i.e. the product of the allele frequency and DoG, is greatest (D). These two scales capture the small scale variation in allele frequency between areas where different alleles are fixed (B), and the large scale variation between the center of the landscape where the alternate allele is present in some locations versus the edges of the landscape where the alternate allele is totally absent (C).

**Fig. S2.**
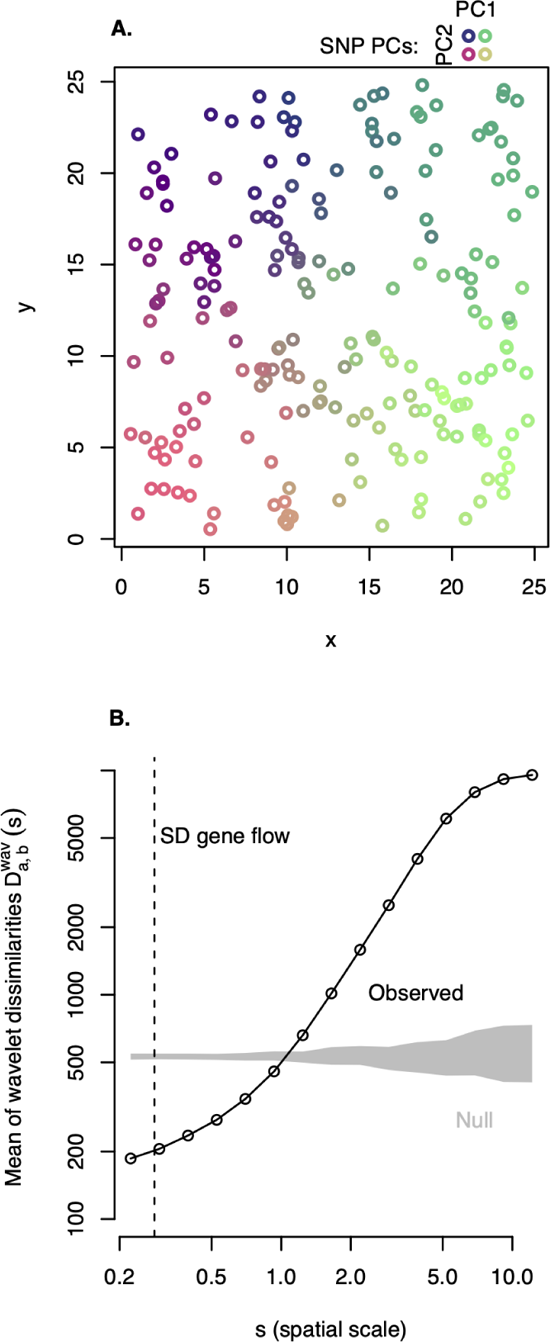
Simulated two dimensional landscape. (A) with continuous population structure among 200 sampled individuals (circles), illustrated by the first two PCs of 1000 randomly selected SNPs (colors). In (A), each individual’s color gives its SNP loadings on PC1 and PC2 according to the key at upper right. Mean of observed wavelet dissimilarities (B) among the 200 samples at a range of spatial scales *s* (connected by a solid black line) in comparison with the null expectation (gray ribbon) from permuted sample locations (2.5-97.5th percentiles of 100 permutations). The standard deviation of gene flow distance is indicated (dashed line).

**Fig. S3.**
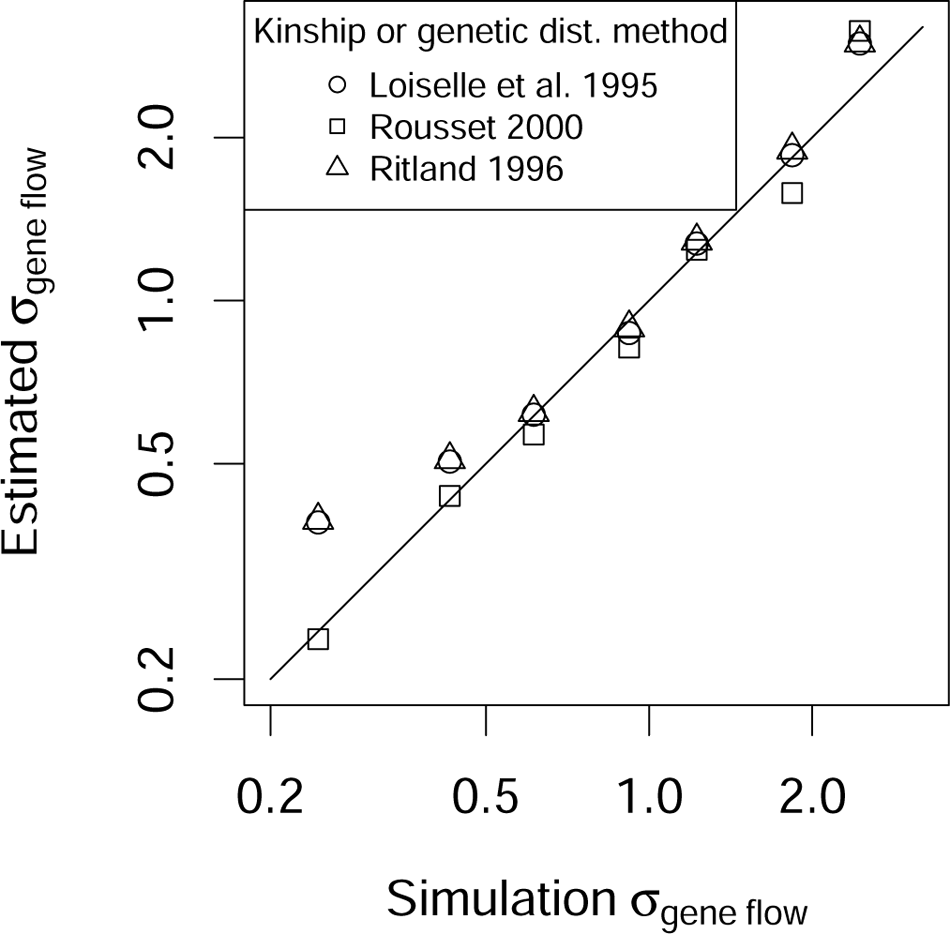
Comparing simulation gene flow parameters (x-axis) with estimations based on theory (y-axis). Three different methods were used to calculate either genetic distance or kinship on 200 individuals and 1,000 random SNPs. The software of Olivier J. Hardy and Xavier Vekemans (2002) was used to estimate gene flow *σ*.

**Fig. S4.**
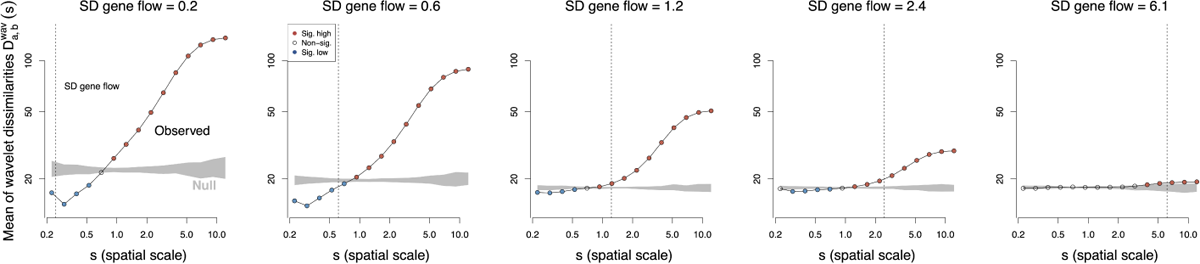
Mean wavelet dissimilarities for a neutral two dimensional landscape with different levels of gene flow in simulation from each panel. 200 samples were taken from each simulation, and wavelet dissimilarities were calculated at a range of spatial scales *s* (connected by a solid black line) in comparison with the null expectation (gray ribbon) from permuted sample locations (2.5-97.5th percentiles of 100 permutations.

**Fig. S5.**
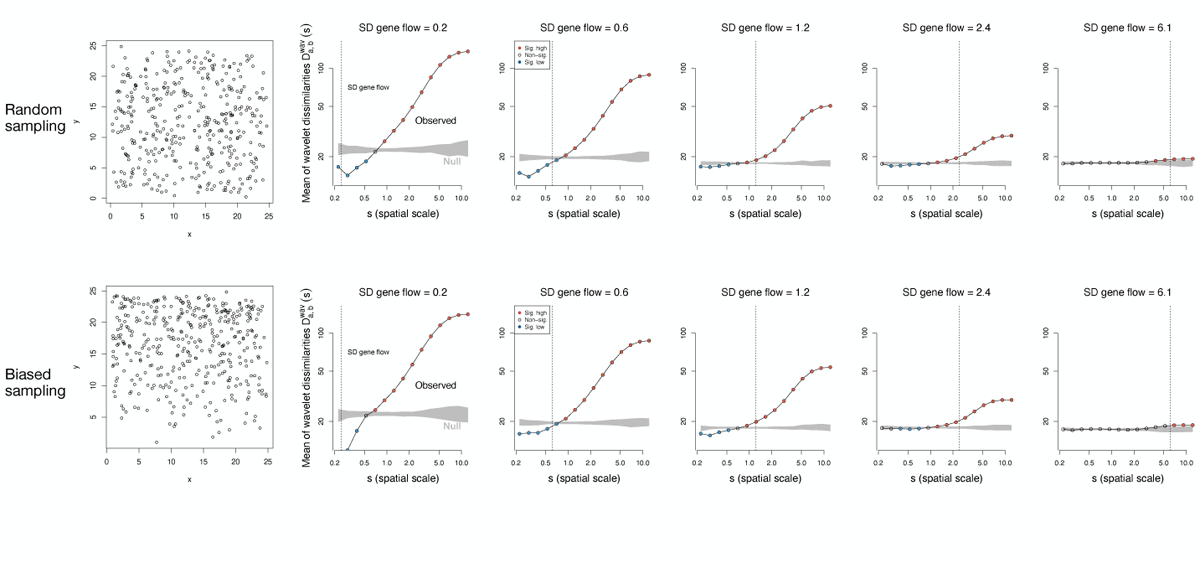
The effect of biased sampling on mean wavelet dissimilarities for a neutral two dimensional landscape with different levels of gene flow in simulation from each panel. 200 samples were taken from each simulation, but in the bottom row these samples were taken so that approximately 3/4 of samples came from the upper 1/2 of the landscape. The comparison with random sampling is shown in the top row.

**Fig. S6.**
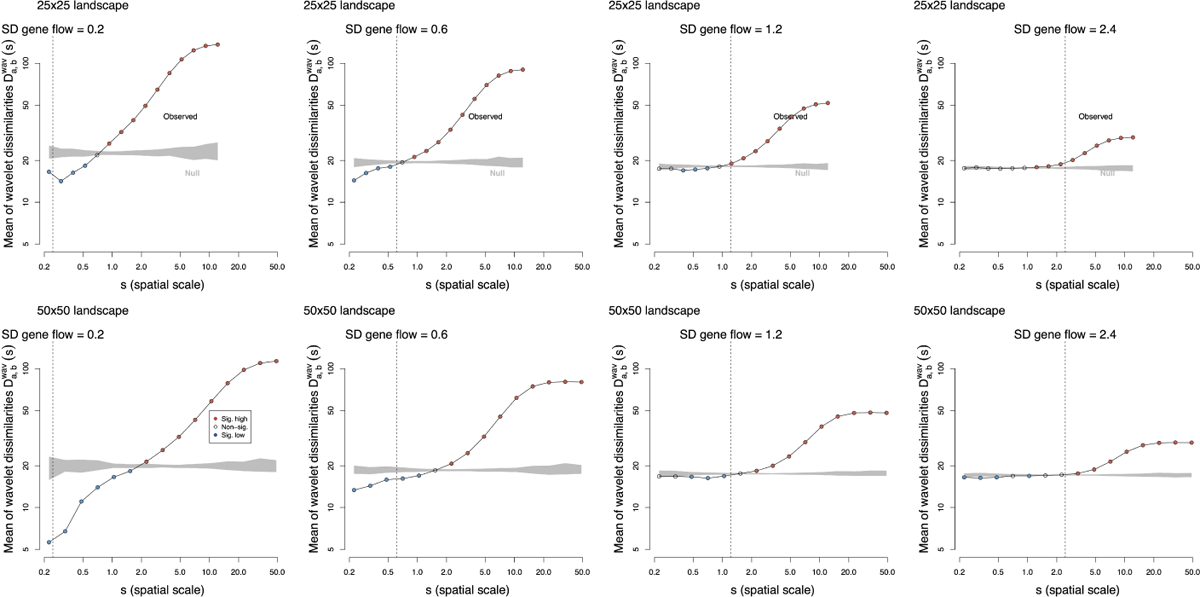
Mean wavelet dissimilarities for neutral two dimensional landscapes of different sizes (25×25 or 50×50) with different levels of gene flow in simulations across each column. 200 samples were taken from each simulation, and wavelet dissimilarities were calculated at a range of spatial scales *s* (connected by a solid black line) in comparison with the null expectation (gray ribbon) from permuted sample locations (2.5-97.5th percentiles of 100 permutations.

**Fig. S7.**
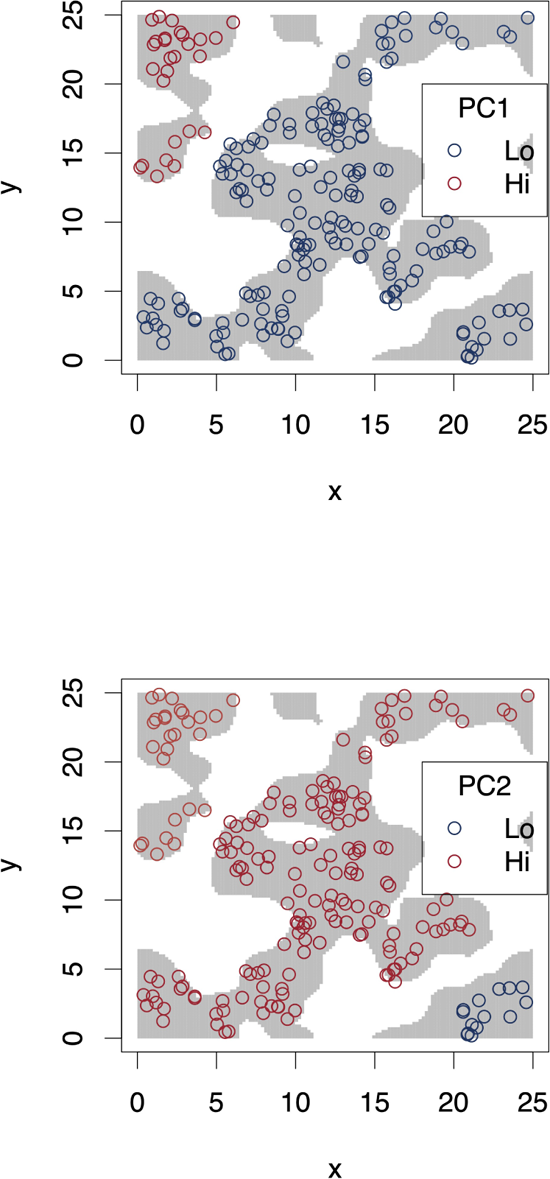
Principal component analysis on 1000 random SNPs from the neutral simulation on a heterogeneous landscape. Habitat is shown as gray in the background and unsuitable areas are white. Sampled individuals are circles. Colors represent the first two PCs and show how the two populations on islands in upper left and bottom right are genetically distinct.

**Fig. S8.**
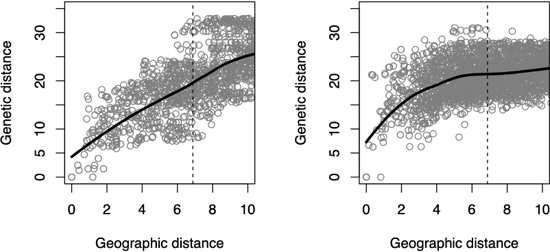
Genetic distance versus geographic distance for 2 regions highlighted in. Figure 3. The two regions highlighted in Figure 3B (gray boxes) are shown here, so that the left panel here corresponds to Figure 3D and the right panel to Figure 3E (the distance in PC1 space is shown in those panels of Figure 3).

**Fig. S9.**
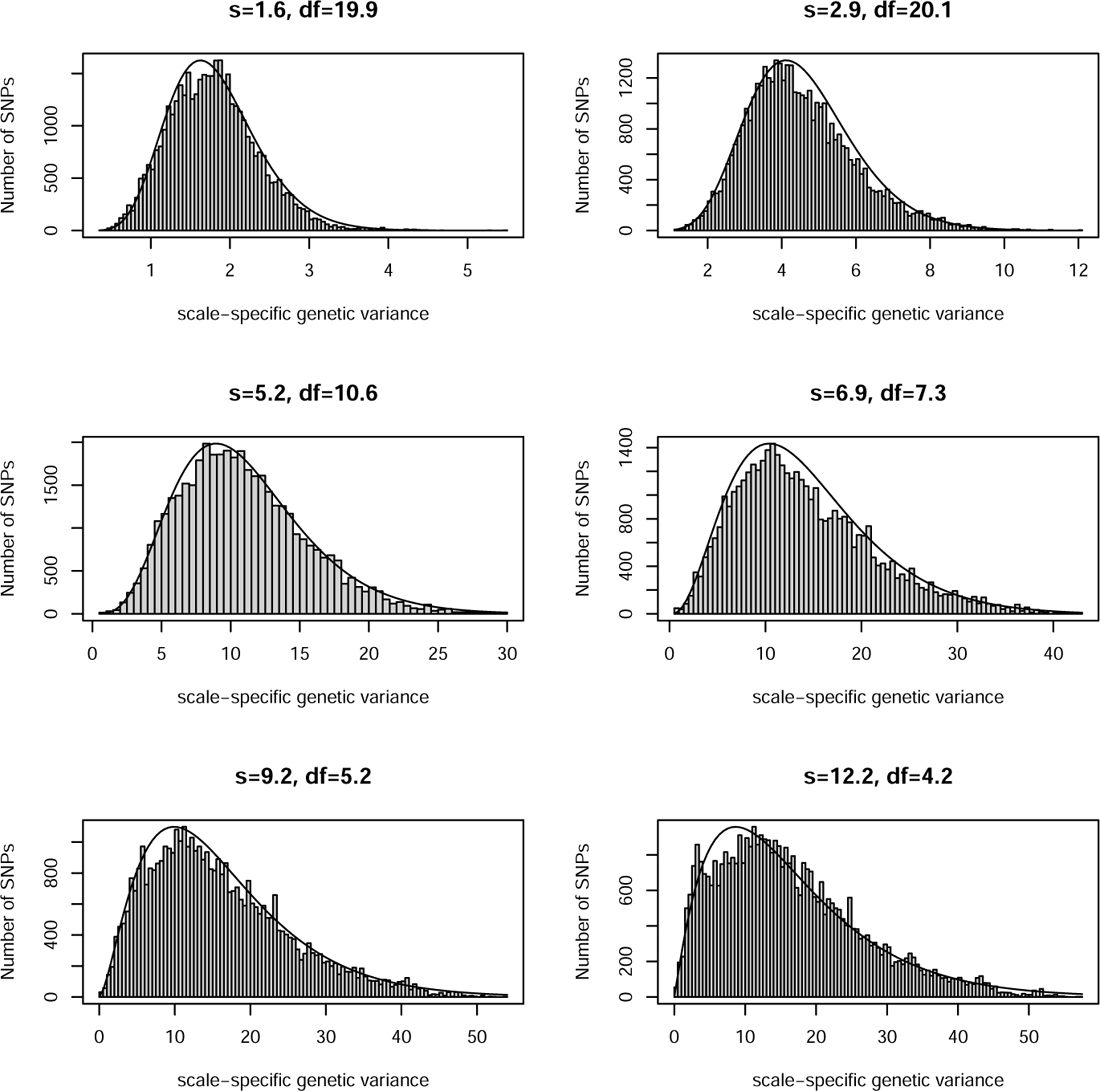
Distribution of scale-specific genetic variance under a neutral scenario after 100,000 time steps compared to fitted. *χ*^2^. Histogram shows SNPs and the curve shows the fitted *χ*^2^. Here the simulation is as in Figures S2 and S3, with *σ* of mating and of dispersal equal to 0.02, for a SD of gene flow equal to 0.24. The degrees of freedom of the *χ*^2^ were fit as described in the main text.

**Fig. S10.**
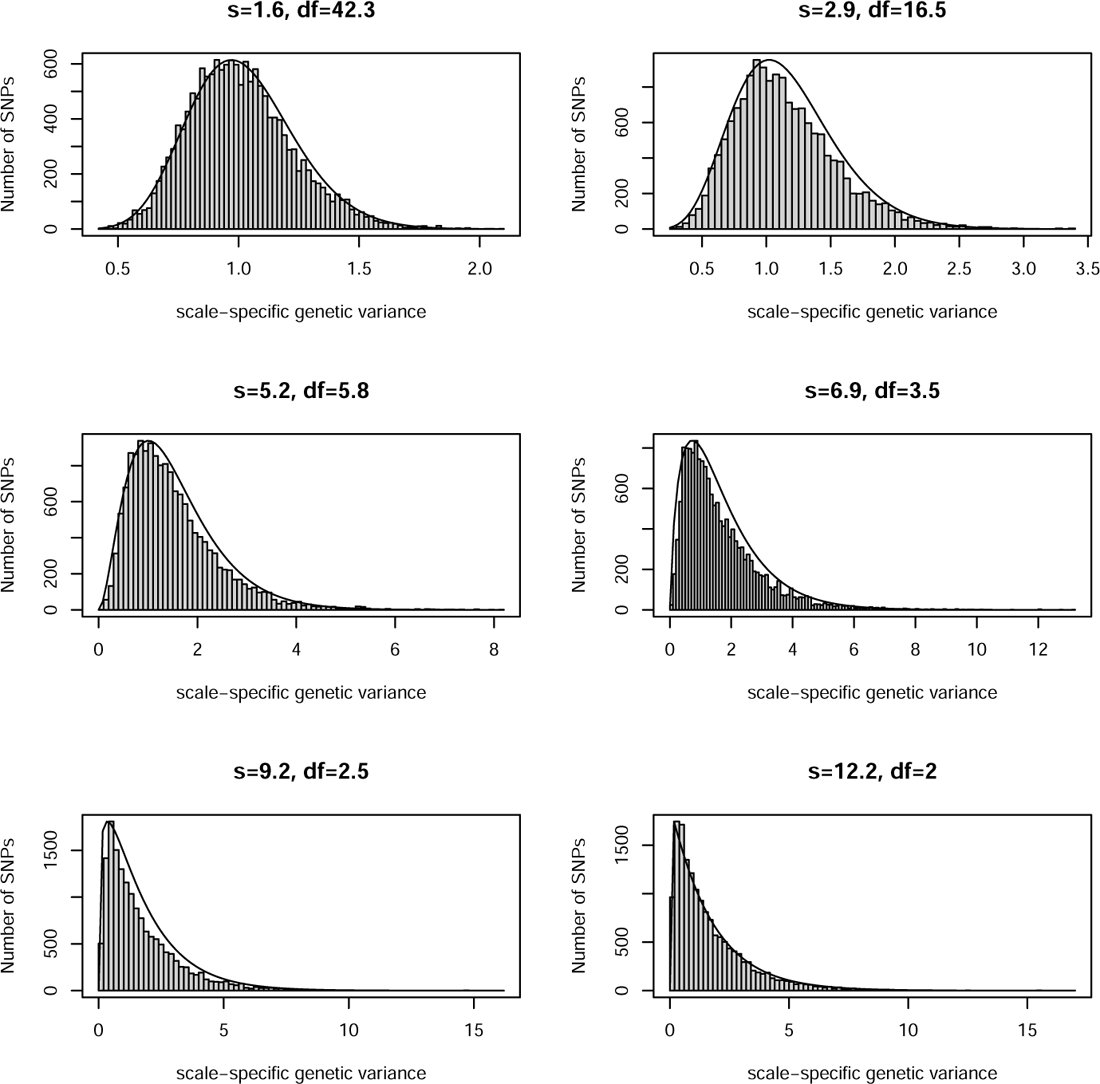
Distribution of scale-specific genetic variance under a neutral scenario after 100,000 time steps compared to fitted. *χ*^2^. Histogram shows SNPs and the curve shows the fitted *χ*^2^. Here the simulation is as in Figures S2 and S3, with *σ* of mating and of dispersal equal to 2, for a SD of gene flow equal to 2.45. The degrees of freedom of the *χ*^2^ were fit as described in the main text.

**Fig. S11.**
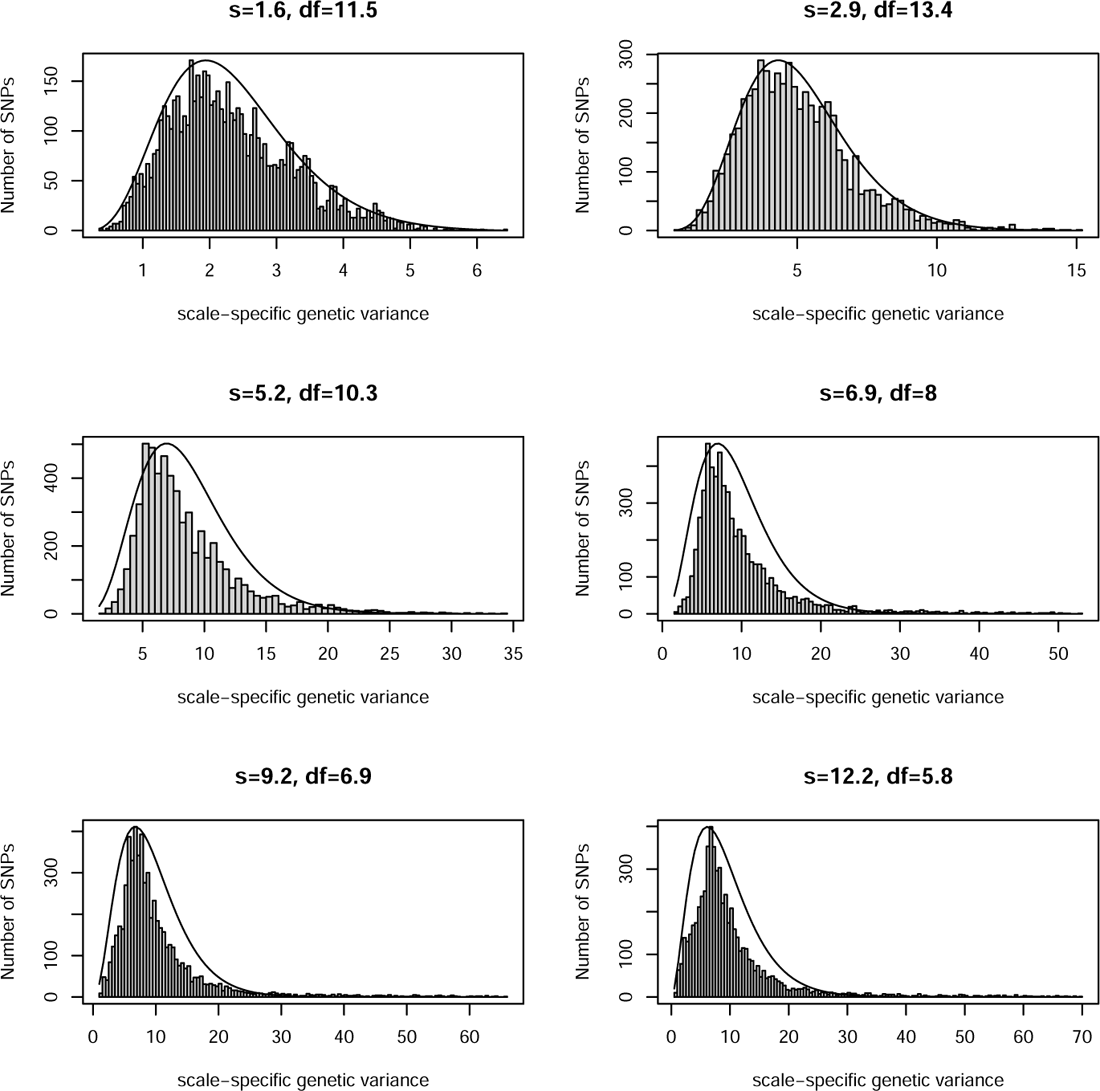
Distribution of scale-specific genetic variance under a neutral scenario with recent rapid range expansion, after 3,000 time steps, compared to fitted *χ*^2^. Histogram shows SNPs and the curve shows the fitted *χ*^2^. Here the simulation is as in Figure 3. The degrees of freedom of the *χ*^2^ were fit as described in the main text.

**Fig. S12.**
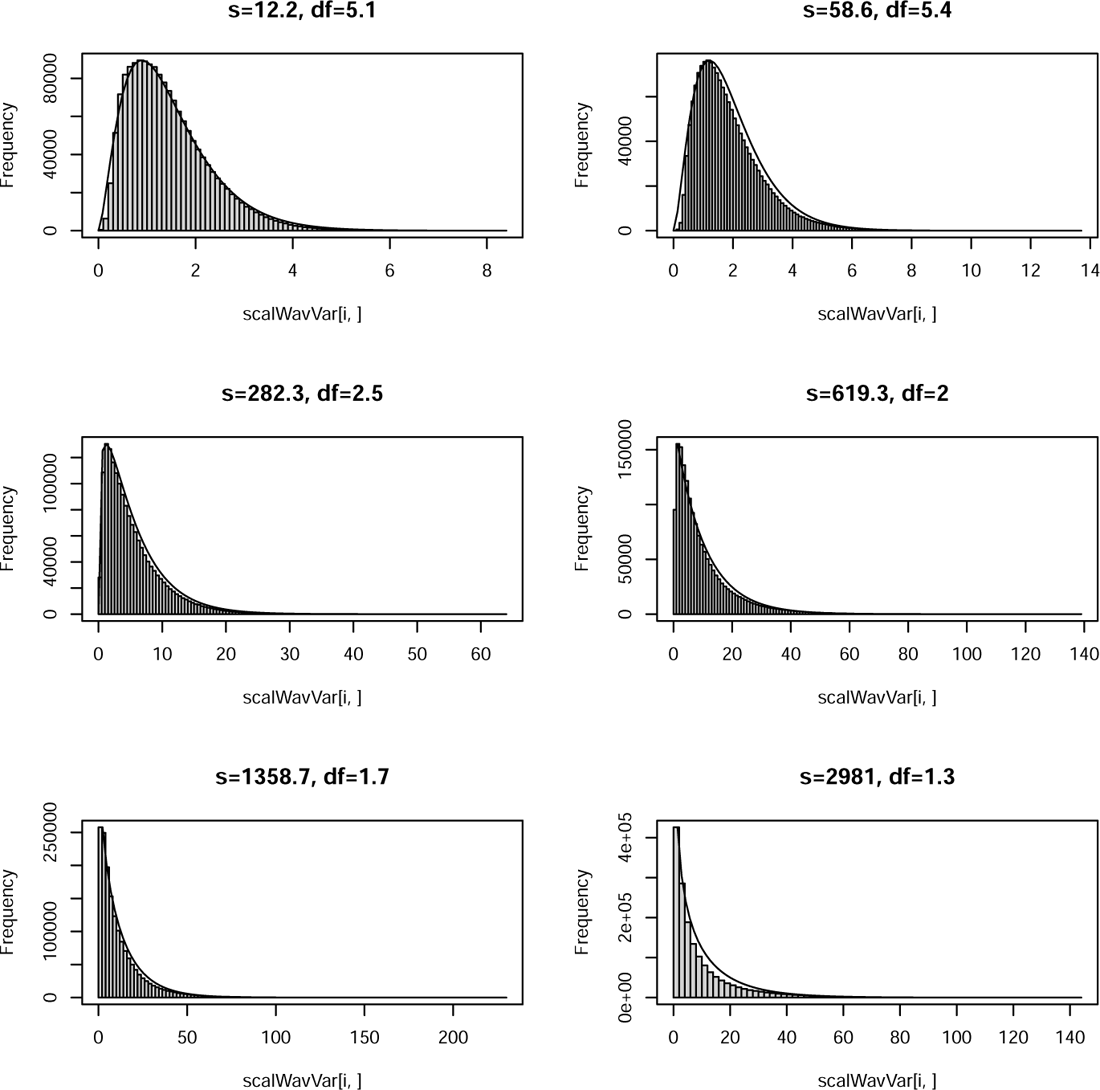
Distribution of scale-specific genetic variance for Arabidopsis, compared to fitted. *χ*^2^. Histogram shows SNPs and the curve shows the fitted *χ*^2^. The degrees of freedom of the *χ*^2^ were fit as described in the main text.

**Fig. S13.**
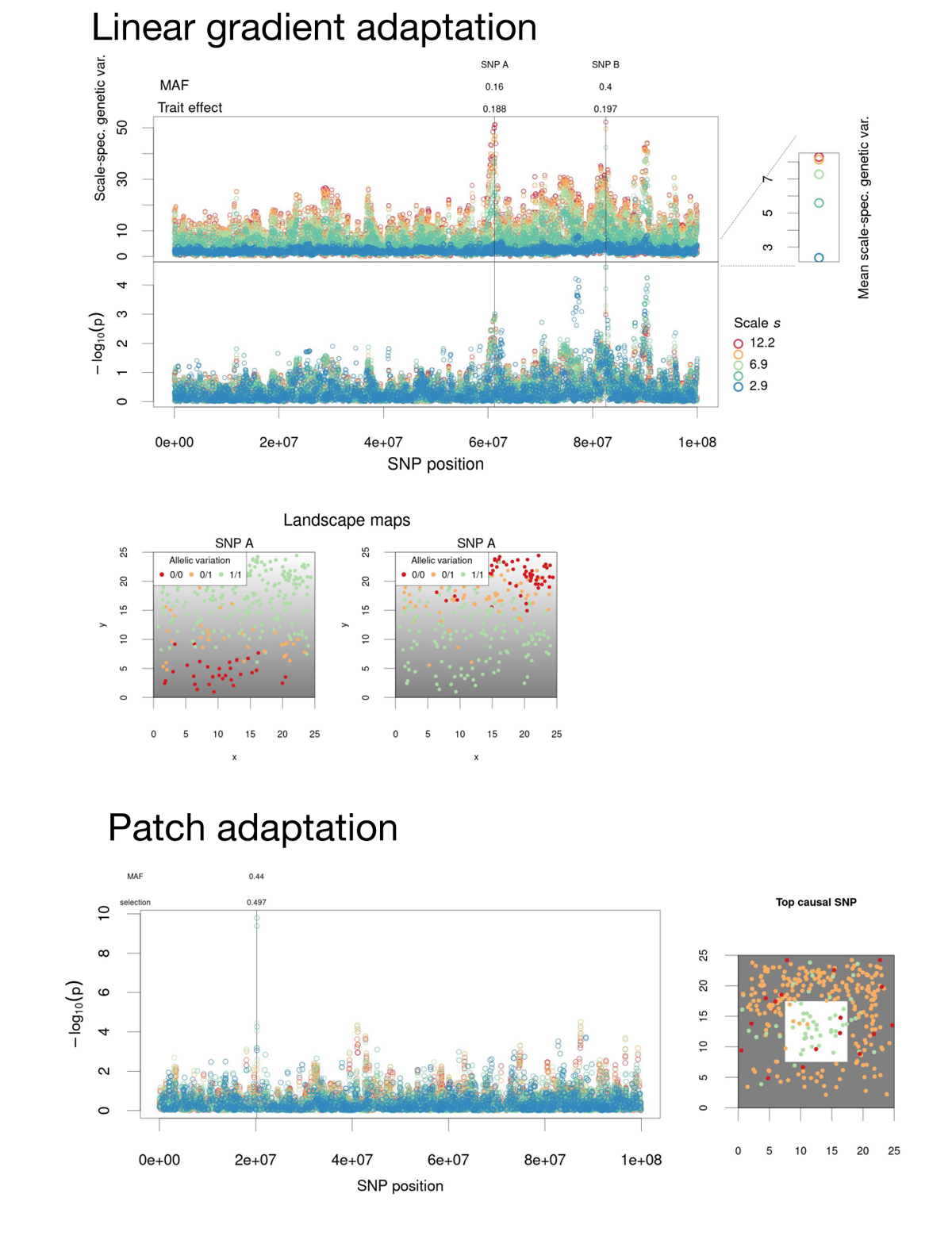
Applying scaled wavelet variance test to two scenarios of local adaptation with heavily biased sampling. These are the same scenarios depicted in Figures 4 and 5 of main text but with heavily biased sampling along the landscape’s y-axis, such that 3/4 of samples are from the upper 1/2 of the landscape, approximately.

**Fig. S14.**
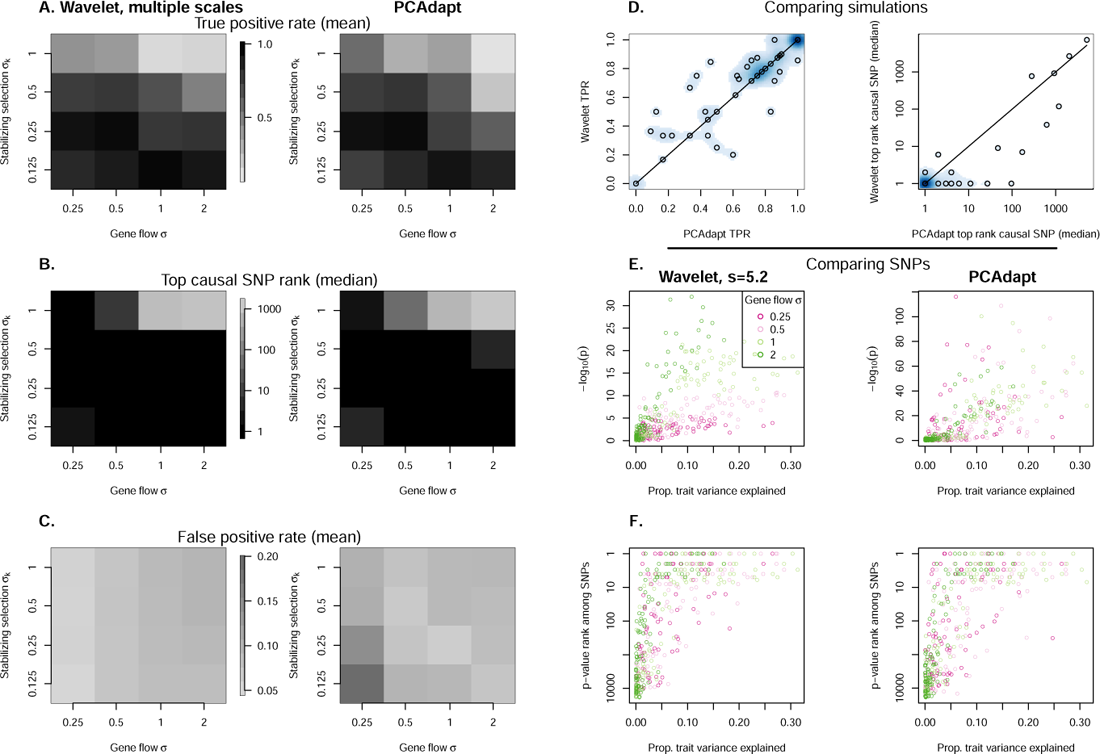
Comparing the scale-specific genetic variance test with PCAdapt in simulations of adaptation to a linear environmental gradient. (A) True positive rates (nominal *p <* 0.05) for each combination of simulation parameters, the scales of mating and dispersal *σ* and the standard deviation of the Gaussian stabilizing selection function *σ_k_*. (B) An alternate view of statistical power based on the median rank of the top selected SNP among all SNPs. (C) False positive rates (nominal *p <* 0.05). (D) Comparing power between the two statistical approaches for the different simulation runs. Density of points is shown in the blue scale so as to indicate where many simulations had the same result. The line indiciates a 1:1 relationship. (E-F) Individual selected SNPs in simulations, showing their nominal *p* values and ranks among all SNPs, colored based on *σ* in the simulation. The x-axis represents the proportion of total phenotypic variation among sampled individuals that was explained by the given SNP (*R*^2^ from a linear model).

**Fig. S15.**
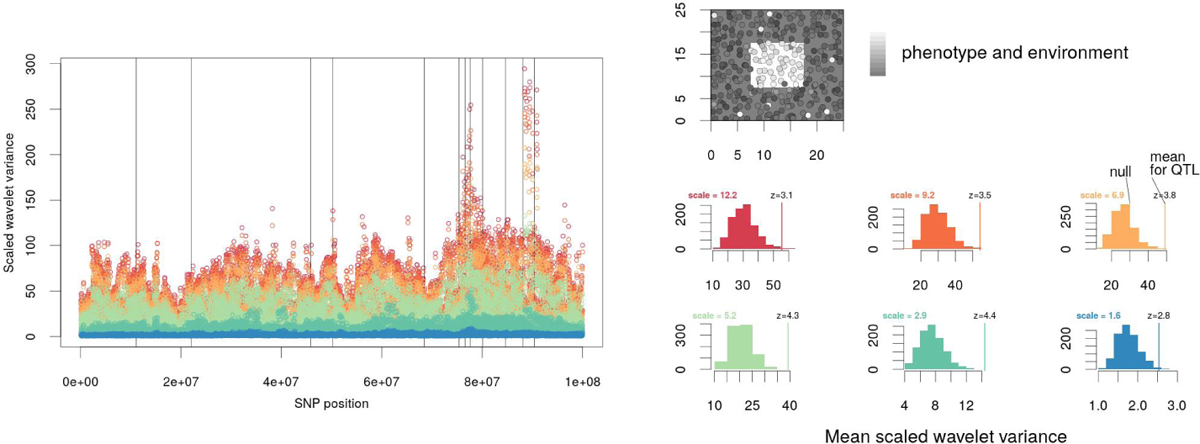
Testing for selection on QTL using wavelet transforms. Comparing mean scale-specific genetic variance *var*((*T^wav^f_i_*)(*a, b, s*)*/sd*(*f_i_*)) for QTL to that of random SNPs, for six different scales *s* (red = large scale and blue = small scale). Populations were locally adapted to a discrete habitat patch and results are shown after 1000 simulated time steps. QTL with MAF of at least 0.05 are indicated with vertical lines at left. The histograms at right show null distributions of mean scale-specific genetic variance *var*((*T^wav^f_i_*)(*a, b, s*)*/sd*(*f_i_*)) based on random samples of an equal number of markers as there were QTL (with MAF at least 0.05, n=13 here) and the observed scale-specific genetic variance of QTL and its z-score.

**Fig. S16.**
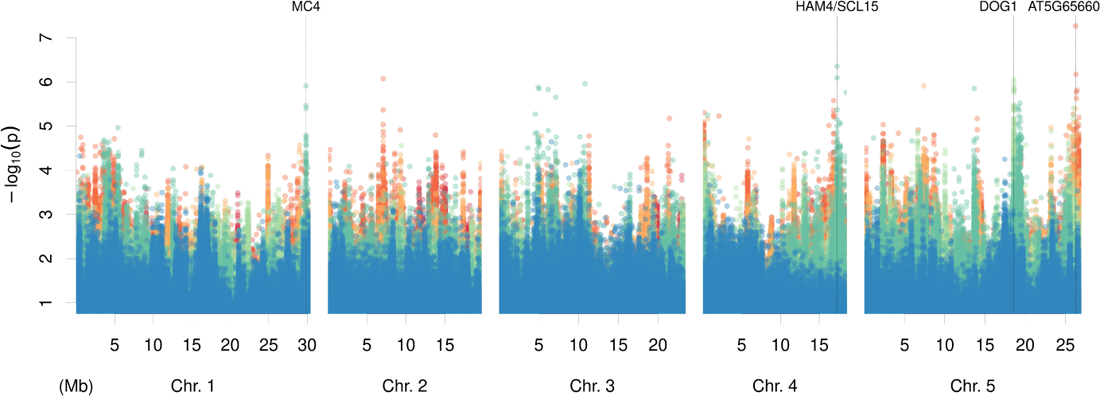
Scale-specific genetic variance test results for SNPs of Arabidopsis. Scales shown go from blue (small scale) to red (large scale), specifically, *∼* 12, *∼* 59, *∼* 282, *∼* 619, *∼* 1359, *∼* 2980 km. Candidate genes are noted.

